# Adenosine signaling in glia modulates metabolic state-dependent behavior in *Drosophila*

**DOI:** 10.1101/2024.08.07.606811

**Authors:** Jean-François De Backer, Thomas Karges, Julia Papst, Cristina Coman, Robert Ahrends, Yanjun Xu, Cristina García-Cáceres, Ilona C. Grunwald Kadow

## Abstract

An animal’s metabolic state strongly influences its behavior. Hungry animals prioritize food seeking and feeding behaviors, while sated animals suppress these behaviors to engage in other activities. Additionally, neuronal activity and synaptic transmission are among the most energy expensive processes. Yet neurons do not uptake nutrients from the circulation. Instead, glia fulfill this highly evolutionary conserved function. Recent studies have shown that glia can modulate neuronal activity and behavior. However, how different glia subtypes sense metabolic state and modulate neurons and behavior is incompletely understood. Here, we unravel two types of glia-mediated modulation of metabolic state-dependent behavior. In food-deprived flies, astrocyte-like and perineurial glia promote foraging and feeding, respectively, while cortex glia suppress these behaviors. We further show that adenosine and adenosine receptor modulate intracellular calcium levels in these glia subtypes, which ultimately controls behavior. This study reveals a new mechanism how different glia subtypes sense the metabolic state of the animal and modulate its behavior accordingly.

## Introduction

Most organisms live in frequently changing environments where nutrient availability fluctuates. Animals must therefore adapt their behavior to food abundance or scarcity and prepare for periods of food deprivation. In order to respond to such conditions, animals adapt their metabolism to mobilize internal energy stores and prioritize behaviors such as food-foraging over other drives^1–3^. However, active search for a food source, food consumption and digestion are themselves energy demanding activities. Those behaviors must therefore be tightly regulated and repressed when the animal reaches satiation. While it is well known that food-related neuronal processing is modulated by metabolic state^4,5^, the role of the activity of the other ubiquitous cell type in the nervous system, glia, in controlling metabolism-related behaviors such as foraging or feeding requires additional investigation.

Neuron-glia communication is starting to emerge as an important regulatory level that shapes behavior^6^. In addition to their long-known role as support cells for neurons, glia, in particular astrocytes, respond to the release of various neurotransmitters, mostly through changes in their calcium levels^7^. In turn, glia modulate neuronal activity via different mechanisms including the regulation of neurotransmitter turn-over, potassium buffering and the release of various signaling molecules often referred to as gliotransmitters^6,8^. Several recent studies have shown that this bi-directional neuron-glia communication shapes numerous behaviors in various model organisms ranging from the nematode *Caenorhabditis elegans* to rodents^6^. Specifically, in the context of metabolic state and obesity, astrocytes regulate neuronal activity in the hypothalamus as well as food intake in mice^9–12^. Given the ubiquitous presence of glia throughout the nervous system as well as their extended arborization and nets, those regulatory mechanisms can occur at the local synaptic level or affecting entire neuronal networks making them ideal candidates for broadcasting essential physiological states globally across the nervous system^8^.

Similar to mammals, glia in insects also comprise different subtypes distributed according to their function within the CNS^13–15^. Cortex glia (CG) tightly embed neuronal somas, which are, contrary to vertebrates, located at the surface of the fly’s brain. There, CG modulate neuronal excitability by potassium buffering^16,17^ and provide necessary nutrients for neurons to sustain memory formation^18^. From the surface, flies’ monopolar neurons send neurites that further differentiate into axons and dendrites, forming the neuropil. Within the neuropil, astrocyte-like glia (ALG) interact with synapses, while ensheating glia (ENG) interact with neurites and forms internal barriers separating neuropil compartments. Several studies have shown that ALG are involved in the modulation of different behaviors in flies, including sleep homeostasis, chemotaxis or drinking^19–21^, while EGN participate in the transmission of negative stimuli during memory formation and are involved in sleep homeostasis^22,23^.

Besides the aforementioned subtypes, two additional glial populations are found in the fly’s brain: the perineurial glia (PNG) and subperineurial (SPN) glia. These glial cells separate the brain and the ventral nerve cord from the circulating hemolymph and function analogous to the mammalian blood-brain-barrier (BBB). Interestingly, despite a variation in ontology, the presence of a barrier isolating the brain from the circulation is remarkably well conserved across animal phyla^24^. In mammals, the BBB is formed by the neurovascular unit that comprises endothelial cells, pericytes, the end-feet of astrocytes, neurons and a basement membrane. By contrast, in flies, the functional compartmentalization between the brain and the circulation is ensured by the PNG and SPN, which form two distinct layers. SPN cells form the layer directly in contact with the nervous tissue containing septate junctions that prevent paracellular diffusion between the hemolymph and the CNS. In addition, SPN cells are crucial for the regulation of nutrient flow by expressing various molecular transporters^25^. Above the SPN, PNG form the outermost layer of the CNS. Like the SPN, they also, contribute to nutrient transport, and are critical to provide energy to the CNS^26^.

Given their position at the interface between circulation and neurons, glia and the BBB have recently been proposed to detect the availability of nutrients in the blood or the hemolymph and to transmit this information to the rest of nervous system^11,27–31^. The mechanisms enabling glia to sense metabolic state are, however, currently unknown. As ATP is the energetic unit of living organisms and can be released and detected in the nervous system by both neurons and glial cells, ATP is a good candidate to bridge metabolic sensing and adaptative behavior in response to high energy demand or deprivation. Within or outside the cytoplasm, ATP can be hydrolyzed into adenosine^32^. Because it is released in extreme cases of energy deprivation such as hypoxia^33^, as well as during extensive muscular activity, inflammation and organ failures, adenosine is sometimes considered a stress signal^34–37^. One role of extracellular adenosine is to contribute to energetic metabolism by signaling cellular energy demand in different tissues^38^. This function is likely well conserved across animal evolution, including in flies^39^.

In the nervous system, higher concentrations of ATP/adenosine were suggested to promote more demanding cognitive tasks when energy is available and lower concentrations to encourage behaviors related to food seeking^31^. According to this model, purinergic signaling would provide information to the brain about the organism’s low energetic context in order to prioritize food search and saving energy through lowering demanding cognitive activity and resting. However, this appealing model remains to be formally demonstrated. In the present study, we provide experimental evidence for such as model. We show that adenosine accumulates systemically in flies during starvation and can be detected by different sub-populations of glial cells, including the ones forming the hemolymph-brain barrier (HBB). Adenosine signaling differentially modifies the calcium levels in different glia subtypes in a metabolic state-dependent manner and modulates behavior to promote food-seeking and feeding. Those findings provide a new mechanism used by flies, and potentially other animals, to adapt their behavior in response to their systemic energy levels.

## Results

### Circulating adenosine modulates feeding behavior in starved flies

Previous studies have shown that adenosine signaling is involved in the regulation of carbohydrate metabolism in fly larvae: mutant flies with increased systemic adenosine levels show deficiencies in glycogen storage and increased concentrations of circulating glucose^40^. In addition, adenosine is produced during infections in response to an increased demand of nutrients by active immune cells^41,42^. We therefore hypothesized that adenosine could be a systemic signal of energy deprivation in starved flies. To test this, we first analyzed adenosine levels in food-deprived flies using mass spectrometry. We observed that adenosine significantly increased in 24h starved flies as compared to fed flies (Figure 1A) suggesting that food deprivation leads to an increase in systemic adenosine levels.

**Figure 1:**
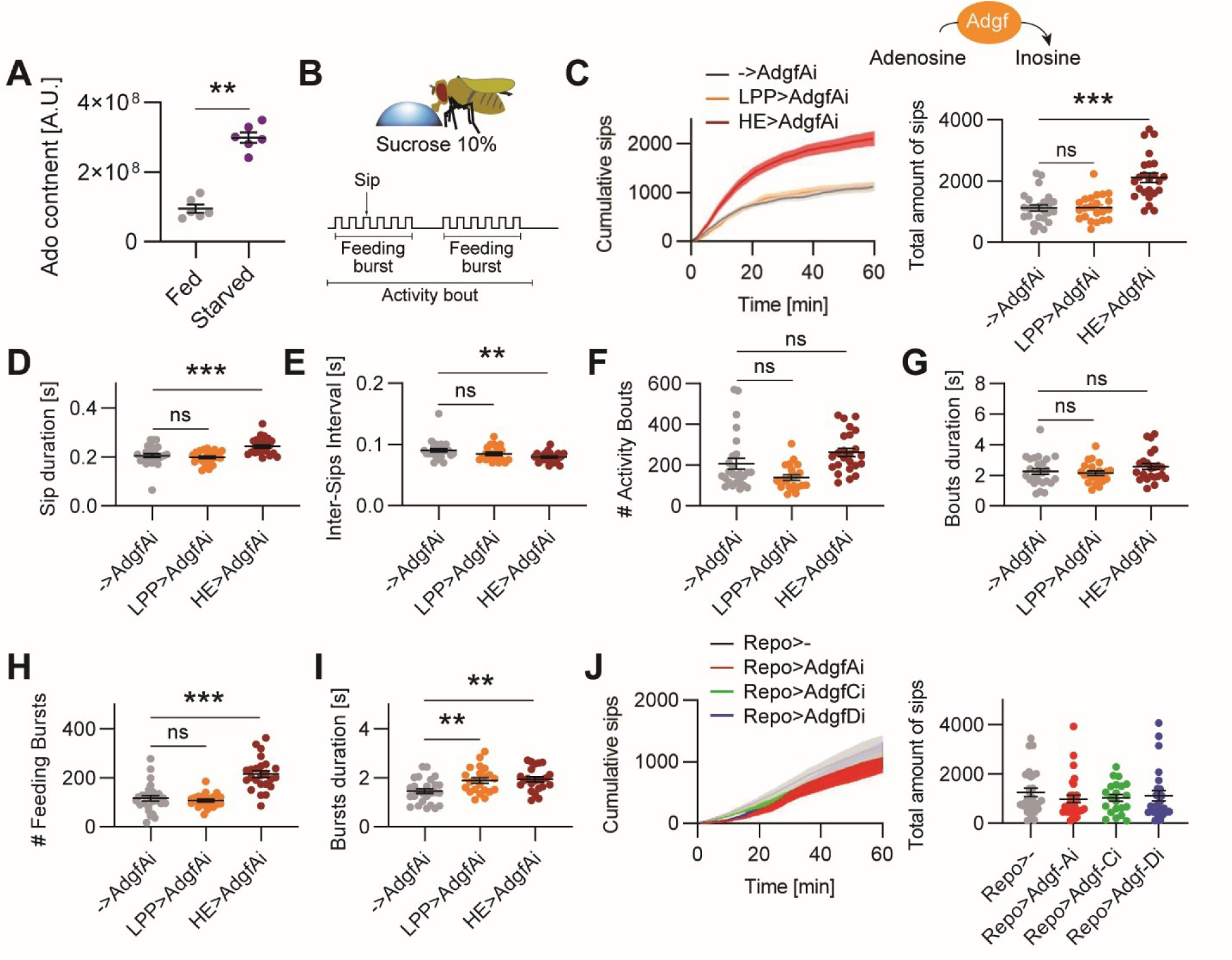
Extracellular adenosine is increased in food-deprived flies and promotes feeding. (**A**) Adenosine content in fed vs. 24h starved CS flies (N=6/6 replicates; 10 flies/replicate; Mann-Whitney U-test p=0.0022). (**B**) Schematic representation of the FlyPAD feeding assay. Single freely-moving flies were feeding on agarose drops containing 10% sucrose during one hour. Feeding behavior was assessed by both the number of sips and the feeding activity pattern. (**C**) In the extracellular space, adenosine is degraded into inactive inosine by adenosine deaminases (Adgf). Cumulative number of sips (mean ± sem) and scatter plot of the total number of sips on 10% sucrose drops in 24h starved control flies (->AdgfAi) or upon knock-down of AdgfA in the fat body (LPP>AdgfAi) or hemocytes (HE>AdgfAi; N=25/22/24; one-way ANOVA p<0.0001). (**D**) Averaged sip duration (one-way ANOVA p<0.0001). (**E**) Averaged time interval between sips (one-way ANOVA p=0.0103). (**F**) Total number of activity bouts (one-way ANOVA p=0.0010). (**G**) Averaged duration of activity bouts (one-way ANOVA p=0.25). (**H**) Total number of feeding bursts (one-way ANOVA p<0.0001). (**I**) Averaged duration of feeding bursts (one-way ANOVA p=0.0017). (**J**) Cumulative number of sips (mean ± sem) and scatter plots of the total number sips taken on 10% sucrose drops in 24h starved control flies (Repo>-) or upon knock-down of the different isoforms of Adgf in glial cells (Repo>AdgfAi, -Ci and -Di; N=27/29/21/25; one-way ANOVA p=0.68). Pair-wise comparisons are indicated as follow: ns, non-significant (p>0.05); *, p<0.05; **, p<0.01; *** p<0.001. See also Figure S1.

Next, we tested if these increased adenosine levels affect feeding-related behavior by manipulating adenosine levels in food-deprived flies. To do this, we knocked down the adenosine deaminase growth factor A (AdgfA), which has previously been shown to increase adenosine levels^43,44^. We performed the knock-down via RNA interference (RNAi) in hemocytes, in which AdgfA promotes the mobilization of internal energy stores^40–42,45^, as well as in the fat body, where the enzyme is not expressed, as a control. To analyze the impact of these manipulations on feeding behavior, we counted the number of sips a single fly was taking during one hour on a drop of agarose containing 10% sucrose using the FlyPAD assay^46^ (Figure 1B; Figure S1A). Interestingly, we found that flies deficient for *AdgfA* in hemocytes took about twice as many sips as the two control strains, flies that only contain the RNAi allele without the driver, as well as flies deficient for *AdgfA* in the fat body (Figure 1C).

In addition to sip count, the temporal resolution of the FlyPAD assay also allows for a precise analysis of the feeding structure^46^. Similar to rodents, flies feed in activity bouts -when the animal visits the food source-which are further subdivided into bursts of licks or sips^46,47^ (Figure 1B).

Consistent with previous findings, 24h food-deprived control flies show an increase in several feeding parameters including sip duration, feeding bursts number and duration as well as the number of activity bouts compared to fed flies^46^ (Figure S1B-G). Knocking-down *AdgfA* in hemocytes of 24h starved flies, and thus elevating the level of circulating adenosine, even further increased several feeding parameters, including sip duration and frequency as well as burst duration and frequency (Figure 1D-I). Taken together, these results suggest that the peripheral production of high levels of circulating adenosine promotes feeding behavior in starved flies.

Next, we asked if adenosine produced by the central nervous system (CNS) could also be involved in the modulation of feeding behavior. The *Drosophila melanogaster* genome contains genes for several members of the Adgf protein family. By analyzing previously published single-cell transcriptomic sequencing data (ssRNAseq) using the SCope online tool^48^, we found that AdgfC and -D are likely expressed in the adult fly brain, mostly in glia. Thus, to assess the role of increased adenosine level in the CNS of starved flies, we knocked-down *AdgfC* and -*D* expression in glia by expressing RNAi under the control of the pan-glia genetic driver Repo-Gal4. As a control, we also expressed the RNAi against *AdgfA*, since the expression of this isoform has not been reported in glia. However, none of these manipulations altered feeding behavior in flies (Figure 1J).

Together, these results suggest that increased levels of adenosine circulating in the hemolymph are responsible for modulating feeding in response to food deprivation.

### Adenosine signaling in the CNS is necessary for feeding state-dependent behavior

Having found that adenosine modulates feeding behavior, we next sought to identify the underlying signaling mechanisms and involved cell types. In contrast to mammals, adenosine signaling in flies is most likely mediated via a single G-protein-coupled adenosine receptor (AdoR)^49^. *AdoR* is preferentially expressed in glial cells, but also in some neurons, as shown by transcriptomic data^48^. To assess if and where AdoR signaling is necessary for metabolic state-dependent behavior, we knocked down *AdoR* in neurons and in glial cells, respectively.

By knocking down *AdoR* in all glia, we observed a significant decrease in the amount of sucrose consumed by 24h starved flies (Figure 2A). Analyzing feeding behavior structure also revealed a reduction in sip duration, sucrose-source visit frequency, and feeding burst frequency and duration (Figure 2B-G), indicating an overall reduction of food consumption. Taken together, these data suggest that systemic adenosine produced during periods of food deprivation stimulates feeding through AdoR signaling in glial cells.

**Figure 2:**
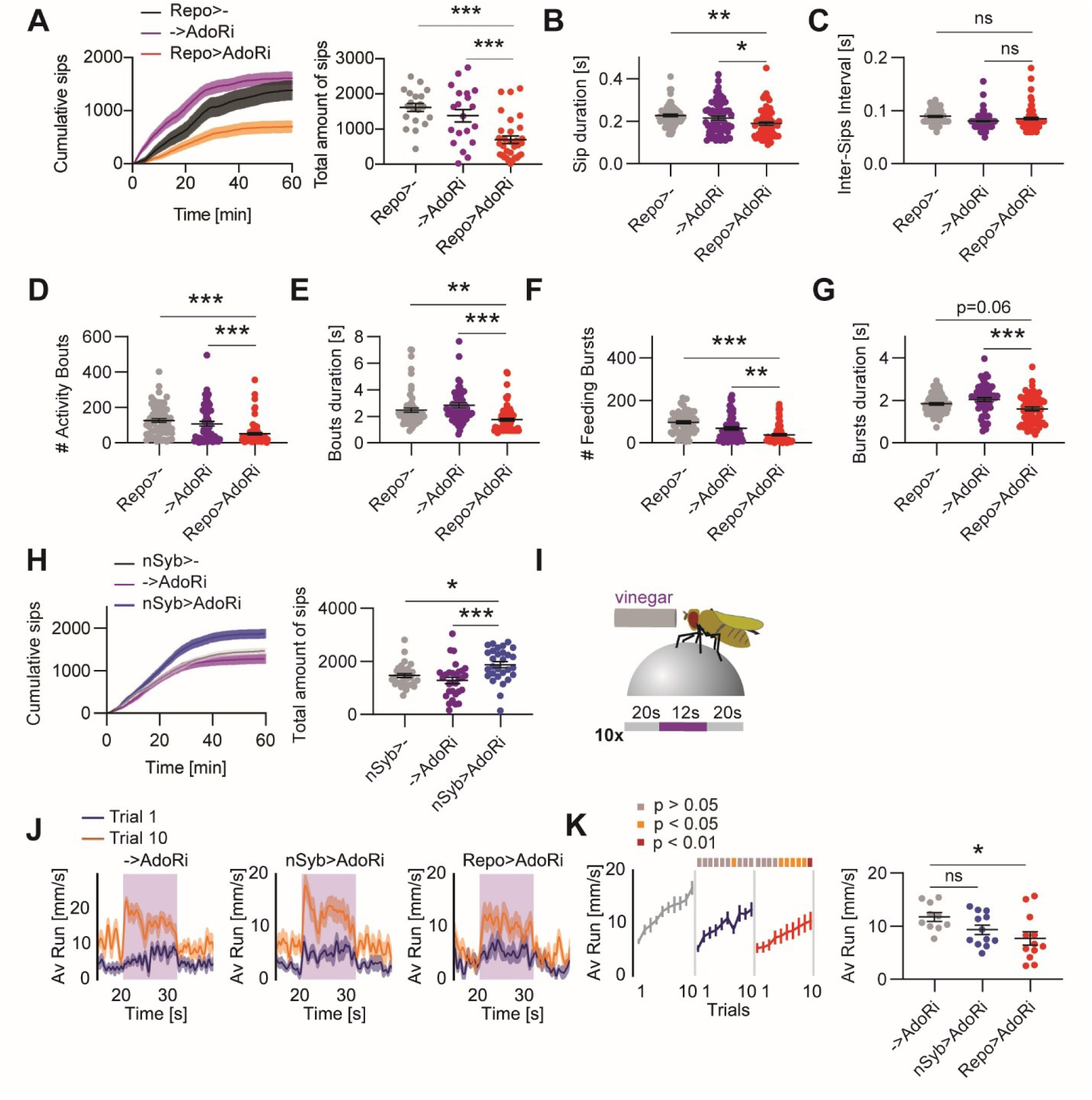
Adenosine signaling in glia is necessary for feeding state-dependent behavior. (**A**) Cumulative number of sips (mean ± sem) and scatter plot of the total number of sips on 10% sucrose drops in 24h starved control flies (Repo>- and ->AdoRi) or upon knock-down of AdoR in glia (Repo>AdoRi; N=20/20/32; one-way ANOVA p<0.0001). (**B**) Averaged sip duration (one-way ANOVA p<0.0001). (**C**) Averaged time interval between sips (one-way ANOVA p=0.0117). (**D**) Total number of activity bouts (one-way ANOVA p<0.0001). (**E**) Averaged duration of activity bouts (one-way ANOVA p<0.0001). (**F**) Total number of feeding bursts (one-way ANOVA p<0.0001). (**G**) Averaged duration of feeding bursts (one-way ANOVA p=0.0008). (**H**) Cumulative number of sips (mean ± sem) and scatter plot of the total number sips on 10% sucrose drops in 24h starved control flies (nSyb>- and ->AdoRi) or upon knock-down of AdoR in neurons (nSyb>AdoRi; N=32/30/29; one-way ANOVA p=0.0005). (**I**) Schematic representation of the odor-tracking paradigm. A single tethered fly is freely walking on an air floating ball and stimulated for 12s with vinegar-odor. This stimulation is repeated over 10 trials (see Methods). (**J**) Averaged forward running speed (± sem) in 24h starved control (->AdoRi) flies or upon knock-down of AdoR in neurons (nSyb>AdoRi) and glia (Repo>AdoRi), respectively. (**K**) Running speed during the stimulus phase over 10 successive trials (mean ± sem; N=10/12/12; 2-way repeated-measures (RM) ANOVA p(groups)=0.0366; p(trials)<0.0001; p(interaction)=0.40). Sidak’s post hoc trial-to-trial comparisons are depicted on the top of the graphs as color-coded boxes (grey p>0.05, orange p<0.05 and red p<0.01). The scatter plot represents the main group effect of the ANOVA. Post hoc pair-wise comparisons are indicated as follow: ns, non-significant; *, p<0.05; **, p<0.01; *** p<0.001. See also Figure S2.

In addition to the FlyPAD feeding assay, we used an olfactory-driven spherical treadmill paradigm developed in our previous work as a proxy for foraging behavior in hungry flies^50^ (Figure 2I). Briefly, we have shown that hungry flies persistently track vinegar odor -a food-predicting cue-with increasing effort over time, while fed flies do not. Here, only the food-deprived flies deficient for *AdoR* in glia, but not in neurons, showed a significant decrease in forward running speed towards the vinegar-odor stimulus (Figure 2J,K), indicating a reduction of the motivation to search for a food-source. We confirmed this result by using another independent RNAi line (Figure S2A,B). To rule out any developmental deficiency caused by the knock-down of *AdoR*, we used the temperature-dependent TARGET system to restrict the expression of the *AdoR*-RNAi to adult flies^51^. Those flies showed a similar decrease in food-odor tracking as those expressing the RNAi through their entire lifespan (Figure S2C-F).

After showing that the knock-down of *AdoR* in glia decreases food consumption, we also tested the role of AdoR in neurons. Surprisingly, flies deficient for *AdoR* in all neurons consumed more sucrose than controls (Figure 2H). Interestingly, *AdoR* seems to be preferentially expressed in dopaminergic (DANs) and octopaminergic neurons^48^ (OANs), two neuron types shown to be key players in the regulation of state-dependent behaviors^5,50^. We therefore knocked down *AdoR* specifically in these two populations of neurons. We found that flies deficient for *AdoR* specifically in DANs, but not in OANs, consumed significantly more sucrose than controls (Figure S2G-M).

Taken together, our data show that the expression of AdoR in glia is not only required for feeding behavior, but also for food-odor tracking in starved flies. The data further suggest that adenosine signaling in DANs could play an additional role in the modulation of different types of behaviors in flies. Moreover, given the very limited current understanding of the role of glia cells in physiological state-dependent behavior, we focused on the role of adenosine signaling in glia in the present study.

### Adenosine signaling in specific glia sub-populations differentially modulates behavior

Using the pan-glia driver line Repo-Gal4, our data indicated that AdoR signaling promotes foraging and feeding in food-deprived flies via glial cells To further dissect the role of adenosine in the different glia subpopulations with different positions and functions in the nervous system, we used subtype-specific driver lines^14^ (Figure 3A).

**Figure 3:**
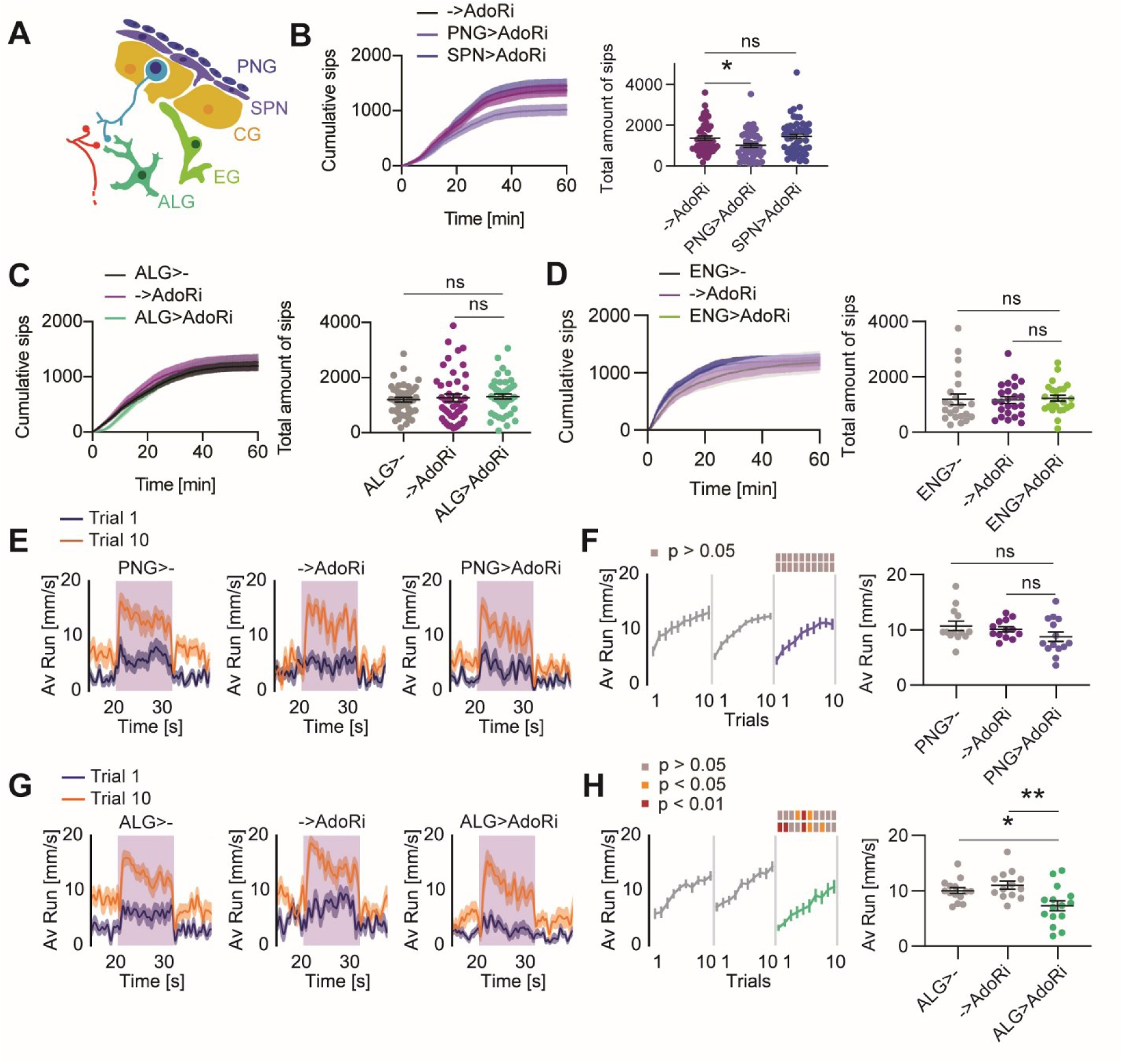
Adenosine signaling in perineurial and astrocyte-like glia subpopulations differentially affect feeding and food-odor tracking behavior. (**A**) Schematic representation of the anatomical location of the different glia subpopulations in the fly CNS. PNG: perineurial glia, SPN: subperineurial glia, CG: cortex glia; ALG: astrocyte-like glia, ENG: ensheating glia. (**B**) Cumulative number of sips (mean ± sem) and scatter plot of the total number sips on 10% sucrose drops in 24h starved control flies (->AdoRi) or upon knock-down of AdoR in PNG (PNG>AdoRi) and SPN (SPN>AdoRi), respectively (N=49/51/48; one-way ANOVA p=0.00118). (**C**) Cumulative number of sips (mean ± sem) and scatter plot of the total number sips on 10% sucrose drops in 24h starved control flies (ALG>- and ->AdoRi) or upon knock-down of AdoR in ALG (ALG>AdoRi; N=47/45/46; one-way ANOVA p=0.72). (**D**) Cumulative number of sips (mean ± sem) and scatter plot of the total number sips on 10% sucrose drops in 24h starved control flies (ENG>- and ->AdoRi) or upon knock-down of AdoR in ENG (ENG>AdoRi; N=22/24/24; one-way ANOVA p=0.95). (**E**) Averaged forward running speed (± sem) in 24h starved control flies (PNG>- and ->AdoRi) or upon knock-down of AdoR in PNG (PNG>AdoRi). (**F**) Running speed during the stimulus phase over 10 successive trials (mean ± sem; N=13/13/14; 2-way RM ANOVA p(groups)=0.19; p(trials)<0.0001; p(interaction)=0.99). (**G**) Averaged forward running speed (± sem) in 24h control flies (ALG>- and - >AdoRi) or upon knock-down of AdoR in ALG (ALG>AdoRi). (**H**) Running speed during the stimulus phase over 10 successive trials (mean ± sem; N=14/13/15; 2-way RM ANOVA p(groups)=0.0030; p(trials)<0.0001; p(interaction)=0.80). For food-odor tracking behavior experiments, Sidak’s post hoc trial-to-trial comparisons are depicted on the top of the graphs as color-coded boxes (grey p>0.05, orange p<0.05 and red p<0.01). The scatter plots represent the main group effect of the ANOVA. Post hoc pair-wise comparisons are indicated as follow: ns, non-significant; *, p<0.05; **, p<0.01; *** p<0.001. See also Figure S3.

Similar to the knock-down of *AdoR* in the entire population of glial cells, 24h starved flies deficient for *AdoR* in PNG took significantly fewer sips of sucrose drops than their genetic control counterparts (Figure 3B). The number of activity bouts and feeding bursts were not significantly different from control flies, while feeding durations were reduced (Figure S3C-F). Contrary to PNG, knock-down of *AdoR* in SPN did not affect the overall amount of consumed food measured by the number of sips in 24h starved flies (Figure 3B). Surprisingly, however, these flies showed a reduced sip frequency and visited the sucrose drops more often than their genetic controls (Figure S3B,C). Nevertheless, this was compensated by shortened durations of feeding activity bouts and bursts, resulting in an overall unaltered food intake (Figure S3D,F). At the level of neuropil glia, we did not observe any change in feeding behavior upon knock-down of *AdoR* in ALG or ENG, respectively (Figure 3C,D). Given that the knock-down of *AdoR* in PNG largely recapitulated the phenotype of pan-glial knock-down of *AdoR*, we conclude that the AdoR expression is specifically required in PNG for an increase in feeding behavior in starved flies.

Contrary to what we observed in feeding behavior, flies deficient for *AdoR* in PNG did not show any alteration of food-odor tracking behavior on the treadmill (Figure 3E,F). Therefore, we analyzed the possible role of AdoR in food-odor tracking in other subtypes of glial cells. Similar to the lack of AdoR in PNG, *AdoR* knock-down in the other surface glia cell-type, the SPN, or in the ENG did not significantly affect vinegar-tracking behavior (Figure S3G-J). By contrast, starved flies deficient for *AdoR* in ALG showed a significant decrease in forward running speed during vinegar-odor presentation (Figure 3G,H) suggesting that AdoR modulates foraging behavior through adenosine signaling in ALG.

Taken together, these data suggest that AdoR is required in distinct subpopulations of glial cells to modulate different aspects of behavior in food-deprived flies. Specifically, AdoR is required in PNG to promote feeding, whereas AdoR signaling in ALG promotes food-odor tracking.

### Adenosine signaling in cortex glia inhibits feeding state-dependent behaviors

So far, we have shown that AdoR signaling promotes foraging and feeding via two different glial subtypes by detecting an increased concentration of adenosine in hungry flies. Interestingly, whereas knock-down of *AdoR* in PNG and ALG recapitulated different aspects of pan-glia knock-down of *AdoR* in feeding and foraging behavior, knock-down of *AdoR* in CG show an opposite phenotype. Indeed, these starved flies took a significantly increased number of sips on the sucrose drops than their genetic controls (Figure 4B). Consistently, they also visited the sucrose source more often compared to control flies and showed an increase in feeding bursts frequency (Figure S4A-F). Re-fed flies (for 40 min on standard food) deficient for *AdoR* in CG showed a trend towards an increase in number of sips, while their genetic controls being undistinguishable from other fed flies (Figure S4H). We did not observe any difference in feeding behavior in these flies while fed (Figure S4G).

**Figure 4:**
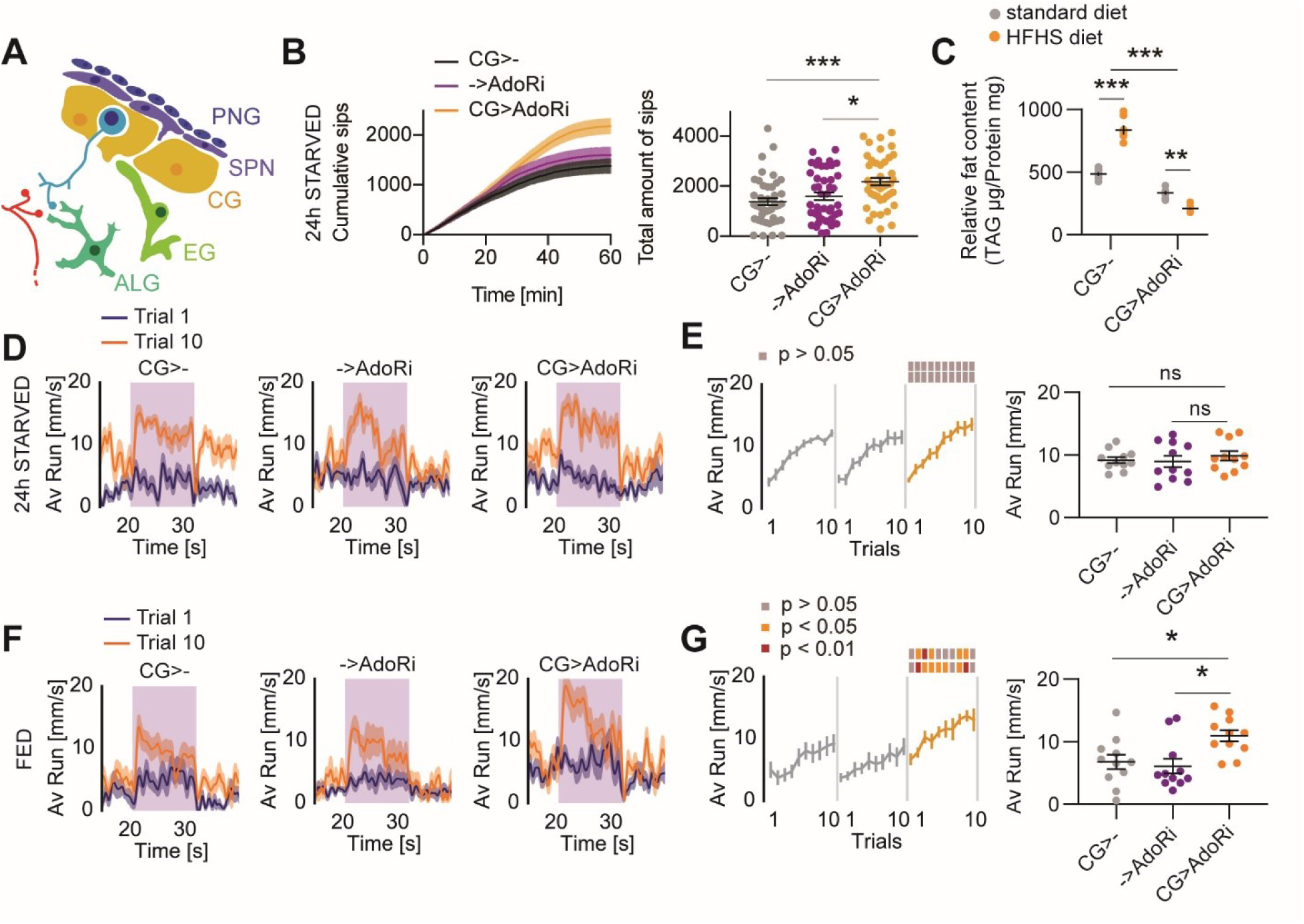
Adenosine signaling in cortex glia inhibits feeding and food-odor tracking behavior. (**A**) Schematic representation of the anatomical location of the different glia subpopulations in the fly CNS. PNG: perineurial glia, SPN: subperineurial glia, CG: cortex glia; ALG: astrocyte-like glia, ENG: ensheating glia. (**B**) Cumulative number of sips (mean ± sem) and scatter plot of the total number sips on 10% sucrose drops in 24h starved control flies (CG>- and ->AdoRi) or upon knock-down of AdoR in CG (CG>AdoRi; N=45/44/44; one-way ANOVA p=0.0004). (**C**) Relative triacylglycerides (TAG) content in control flies (CG>-) and flies deficient for AdoR in CG (CG>AdoRi) fed on standard fly-food or on high-fat, high-sugar containing food (HFHS; N=8/7/8/8 replicates; 5 flies/replicate; 2-way ANOVA p(groups)<0.0001; p(diet)<0.0001; p(interaction)<0.0001). (**D**) Averaged forward running speed (± sem) in 24h starved control flies (CG>- and ->AdoRi) or upon knock-down of AdoR in CG (CG>AdoRi). (**E**) Running speed during the stimulus phase over 10 successive trials (mean ± sem; N=11/11/11; 2-way RM ANOVA p(groups)=0.67; p(trials)<0.0001; p(interaction)=0.40). (**F**) Averaged forward running speed (± sem) in fed control flies (CG>- and ->AdoRi) or upon knock-down of AdoR in CG (CG>AdoRi). (**G**) Running speed during the stimulus phase over 10 successive trials (mean ± sem; N=11/11/11; 2-way RM ANOVA p(groups)=0.0077; p(trials)<0.0001; p(interaction)=0.50). For food-odor tracking behavior experiments, Sidak’s post hoc trial-to-trial comparisons are depicted on the top of the graphs as color-coded boxes (grey p>0.05, orange p<0.05 and red p<0.01). The scatter plots represent the main group effect of the ANOVA. Post hoc pair-wise comparisons are indicated as follow: ns, non-significant; *, p<0.05; **, p<0.01; *** p<0.001. See also Figure S4.

In addition, 24h starved flies deficient for *AdoR* in CG did not show any difference in their running speed during vinegar stimulation in odor tracking experiments (Figure 4D,E), while flies fed *ad libitum* ran significantly faster towards vinegar-odor than controls in successive trials (Figure 4F,G). These data show that appropriate, energy-dependent, expression of foraging and feeding behavior requires expression of AdoR in CG.

Since flies deficient for *AdoR* in CG fail to sense their high energy levels and continue to feed, we wondered about their ability to store superfluous energy in the form of fat. We measured the flies’ triacylglycerides (TAG) lipids levels after being fed on a high fat, high sugar diet, known to induce an accumulation of fat in the fly’s body (Figure 4C). Contrary to control flies, flies with *AdoR* knock-down in CG did not show the expected increase in their TAG/protein ratio. Instead they presented a significantly decreased fat content, as compared to flies fed on standard medium (Figure 4C). This suggests that fat storage is impaired upon loss of *AdoR* in CG, consistent with the interpretation that AdoR in CG is involved in the control of metabolic processes in addition to behavior.

Regarding behavior, our data show that adenosine signaling in glia modulates hunger state-dependent behaviors, i.e. foraging and feeding, differentially through at least three different types of glial cells, PNG, ALG and CG. While AdoR in PNG promotes feeding but not foraging, ALG AdoR signaling promotes foraging but not feeding upon starvation. By contrast, AdoR signaling in CG is required to suppress foraging and feeding in flies fed *ad libitum*.

### Starvation and adenosine signaling modulate glial intra-cellular calcium levels

Our data indicate an important role of adenosine signaling in different glial subpopulations in the regulation of feeding and foraging behavior. We next wanted to understand the mechanisms linking adenosine signaling in glia to our observed phenotypes. In different species, calcium activity in glia may occur in response to neuronal activity and in turn trigger events that modulate neuronal excitability and synaptic transmission, and ultimately, animal behavior^6,8^. In flies, glial calcium activity in relation to behavior and/or neuronal excitability has also been shown^16,17,19,20,22,52,53^. More recently, de Treddern et al.^54^ showed that CG expresses nicotinic cholinergic receptors and that the application of nicotine triggers calcium influx suggesting that these glia might respond to neuronal activity. We therefore investigated the influence of adenosine and AdoR on glial cell function using calcium levels as a proxy for cellular activity.

To analyze whether starvation modulates glial calcium activity, we used the calcium-modulated photoactivable ratiometric integrator (CaMPARI2) and investigated cytoplasmic calcium levels in PNG in explant brains^55,56^. In standard extracellular saline which contains the main types of sugars found in the hemolymph (D-glucose and trehalose), we did not observe any difference in photoconverted CaMPARI2 signals between fed and 24h starved flies (Figure S5A,B). However, the sugar content in hemolymph is known to drop during starvation^57^. We thus repeated this experiment using a saline solution that does not contain either D-glucose or trehalose^58^. Under these conditions, we observed a significant increase in calcium-bound CaMPARI2 in the brain of 24h starved flies as compared to fed flies suggesting that PNG react to low levels of sugar, or starvation, with an increase of cellular calcium (Figure 5A,B). Since AdoR is necessary in PNG for feeding behavior in starved flies, we assessed the cytoplasmic calcium level in flies with a knock-down of *AdoR* in PNG cells. Here, we observed that downregulation of *AdoR* expression indeed prevented calcium increases in brains from 24h starved flies incubated in sugar-free saline, indicating that adenosine signaling is necessary for starvation-induced calcium increase in PNG (Figure 5A,B).

**Figure 5:**
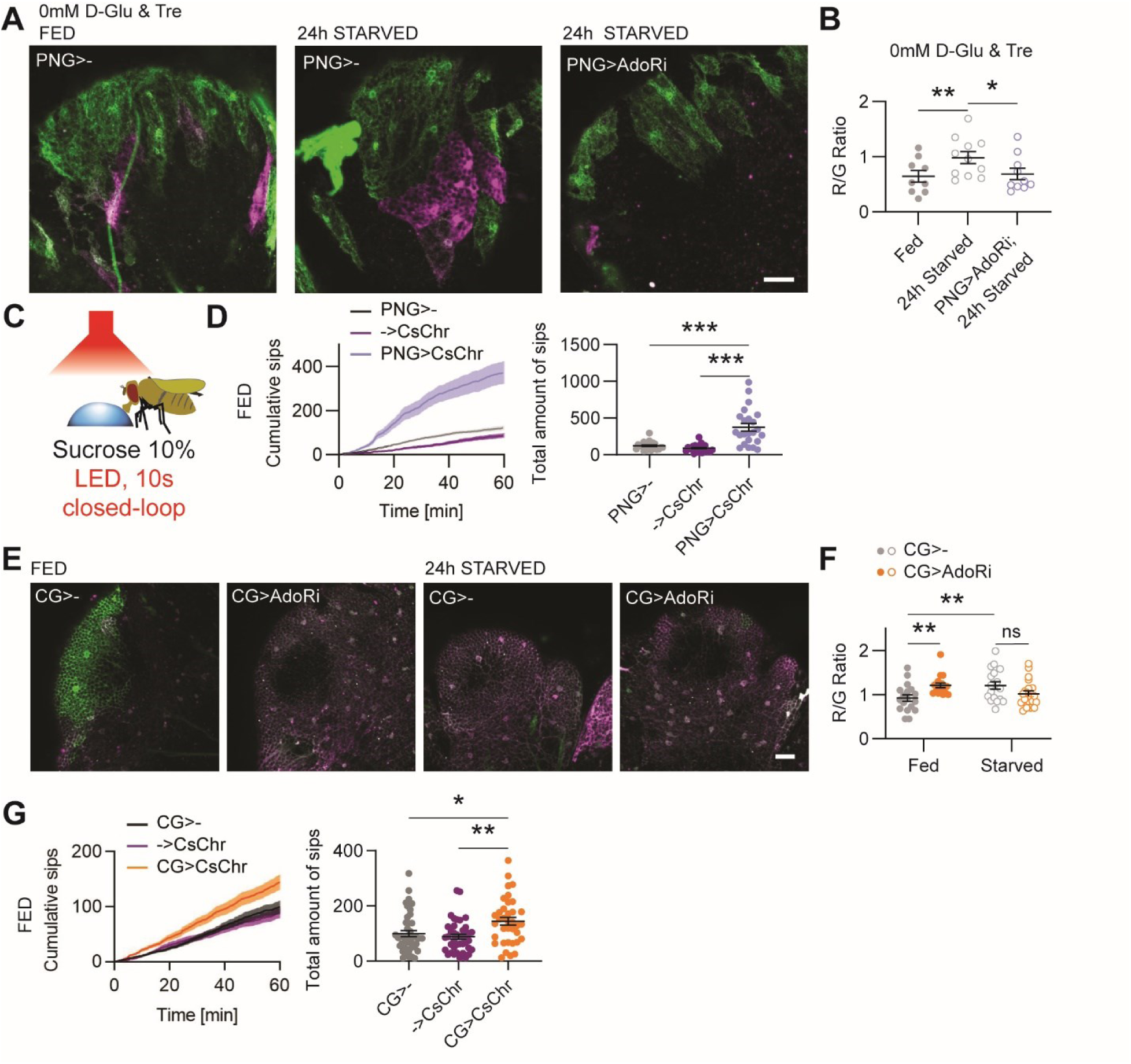
Starvation and adenosine signaling modulate intra-cellular calcium levels in perineurial and cortex glia. (**A**) Example images of CaMPARI2-L398T-expressing PNG cells in dorso-caudal brain explants from fed and 24h starved control flies (PNG>-) and upon knock-down of AdoR (PNG>AdoRi), in sugar-free artificial hemolymph saline (AHL), after photoconversion (D-Glu: D-glucose; Tre: trehalose; scale bar = 20µm). (**B**) Scatter plot of photoconversion ratios in fed and 24h starved control flies (N=9/11), as well as for 24h starved PNG>AdoRi flies (N=10; 2-way ANOVA p(AHL)=0.0076; p(groups)=0.06; p(interaction)=0.06). (**C**) Schematic representation of the closed-loop optogenetic stimulation paradigm used in the FlyPAD feeding assay. The red LED is triggered by the interaction between the fly’s proboscis and the sucrose drop and remains on for 10s. (**D**) Cumulative sips (mean ± sem) and scatter plot of the total sips on 10% sucrose drop in fed control flies (PNG>- and ->CsChr) or in fed flies expressing CsChrimson in PNG (PNG>-; N=23/22/23; one-way ANOVA p<0.0001). (**E**) Example images of CaMPARI2-L398T-expressing CG cells in dorso-caudal brain explants from fed and 24h starved control flies (CG>-) and upon knock-down of AdoR (CG>AdoRi), after photoconversion (scale bar=20µm). (**F**) Scatter plot of photoconversion ratios in fed and 24h starved control flies (CG>-; N=18/17) and upon knock-down of AdoR (CG>AdoRi; N=18/20; 2-way ANOVA p(feeding state)=0.50; p(genotype)=0.49; p(interaction)=0.0014; see Figure S5I). (**G**) Cumulative sips (mean ± sem) and scatter plot of the total sips on 10% sucrose drops in fed control flies (CG>- and ->CsChr) or in fed flies expressing CsChrimson in CG (CG>CsChr; N=44/41/37; one-way ANOVA p=0.0022). Post hoc pair-wise comparisons are indicated as follow: ns, non-significant; *, p<0.05; **, p<0.01; *** p<0.001. See also Figure S5.

In light of these results, we decided to artificially increase the calcium levels in PNG by expressing the red-shifted channelrhodopsin CsChrimson^59^. Although glial cells are not electrically excitable, channelrhodopsins are known to be permeable for calcium cations and have been successfully used in glia to induce calcium transients^60,61^. We optogenetically activated glial cells on the FlyPAD when the fly’s proboscis contacted the drop of sucrose^62^ (Figure 5C). We observed that optogenetic stimulation of PNG during feeding progressively induced an increase in the amount of sucrose consumed by fed flies (Figure 5D). Similar to what we observed in wild-type 24h starved flies, increased calcium activity in PNG was associated with an increase in sip duration, activity bouts and feeding bursts frequencies (Figure S5C-H), suggesting that our optogenetic stimulation paradigm indeed mimicked food deprivation in fed flies.

Next, we measured intracellular calcium levels in CG cells in fed and food-deprived animals. We again observed a higher photoconversion rate of CaMPARI2, indicated higher calcium levels, in brains from 24h starved flies (Figure 5E,F). Importantly, and in contrast to our observations in PNG, knocking-down *AdoR* in these cells increased the calcium concentration in CG of brains from fed flies, but not in starved flies (Figure 5E,F), suggesting that AdoR is necessary to maintain calcium at lower levels in the CG of fed flies. However, although CG is also thought to transport glucose^25,54^, the presence of D-glucose and trehalose in the saline solution did not influence the calcium level in CG in fed or starved brain, respectively (Figure S5I).

We again used CsChrimson to induce higher calcium levels in CG during behavioral experiments. Albeit not with the same amplitude as for PNG, closed-loop optogenetic stimulation of CG also mildly increased the number of sips taken by fed flies on drops of sucrose, correlated with an increase in activity bouts and feeding bursts (Figure 5G; Figure S5J-O). These data, together with the acute stimulation with sugar in the imaging experiments indicate a more complex relationship between sugar, calcium levels, and feeding behavior. Nevertheless, our imaging experiments comparing *AdoR* knock-down in different glia cell types are in line with the *AdoR*-RNAi behavior experiments (see Figure 4) and show that AdoR suppresses both increased calcium levels in CG and increased feeding behavior.

Altogether, our data show that AdoR signaling is responsible for the regulation of cytoplasmic calcium levels in PNG and CG. While adenosine signaling in PNG is necessary for the calcium increase we observed in starved flies, it prevents its starvation-induced increase in CG of fed flies.

### Adenosine signaling in perineurial and cortex glia alters dopaminergic neurons calcium responses in a state-dependent manner

Since it is probable that the regulation of behavior through glial cells is executed through neurons, we next sought to identify candidate neuron types that might contribute to this mechanism. Given the important role of dopamine in the regulation of state-dependent behaviors and prior work in rodents and flies demonstrating that astrocytes modulate DAN activity^19,63–65^, we focused on these modulatory neurons. The fly brain contains several clusters of DANs with prominent neurons innervating the mushroom body (MB). We and others have previously shown that DAN responses to odors are modulated by metabolic state^66,67^. Moreover, we have shown that MB DANs modulate foraging behavior^50^. Specifically, DANs from the PAM cluster innervate the horizontal lobe of the MB and respond to various appetitive signals, including vinegar-odor and sucrose, in a metabolic state-dependent manner^66–69^. We thus assessed the role of glia in the modulation of PAM DAN responses to food-related sensory cues. Using *in vivo* 2-photons microscopy, we recorded calcium activity in PAM neurons in response to vinegar-odor stimulations in flies with and without AdoR in PNG or CG (Figure 6A-C). In control flies, we observed an increased vinegar-odor response in 24h starved flies, as compared to fed flies, consistent with our previous findings^66^(Figure 6D-K). Upon knock-down of *AdoR* in PNG, we observed an increase in vinegar-odor response in most of the compartments of the MB of fed flies but not in starved flies (Figure 6D-K). Only PAM neurons innervating the γ3 compartment did not show this loss of feeding state modulation (Figure 6J,K), suggesting that the change in PNG activity does not affect all neurons homogeneously. Importantly, in flies where *AdoR* was knocked-down in CG, we observed the opposite phenotype as compared to flies with a knock-down of *AdoR* in PNG, just like what we have seen in the behavioral experiments where *AdoR* knock-down in PNG decreased feeding but increased it in CG. The calcium response to vinegar-odor was decreased in starved flies (Figure 6D-K).

**Figure 6:**
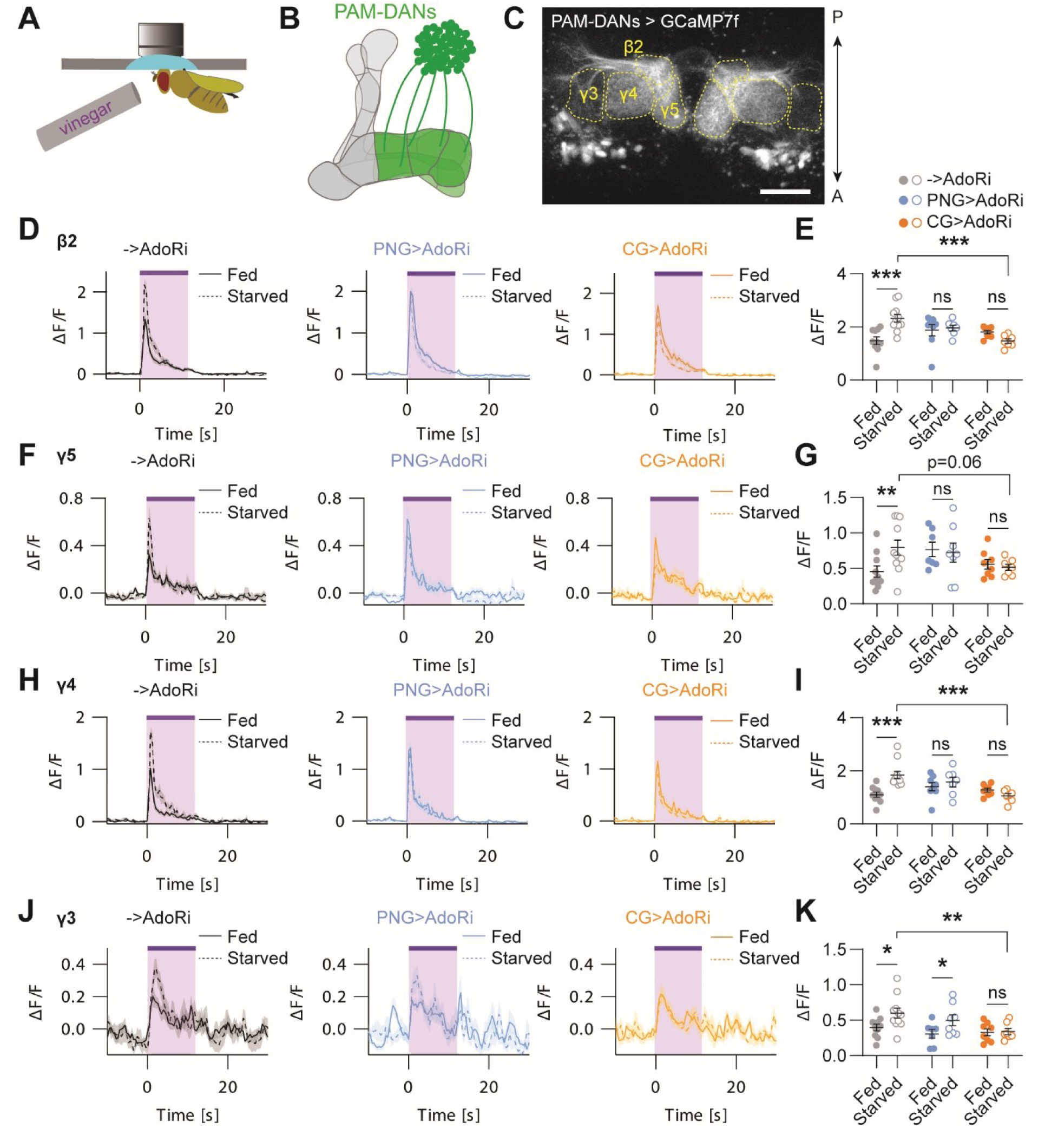
Adenosine signaling in perineurial and cortex glia differentially modulates dopaminergic neurons response to vinegar in fed and starved flies. (**A**) Schematic representation of the *in vivo* imaging set-up. Tethered flies were imaged under a 2-photons microscope and exposed to vinegar-odor stimulation for 12 seconds. (**B**) Schematic representation of the PAM-DA neurons innervating the horizontal lobe of the mushroom body (MB). (**C**) Representative z-averaged projection image showing the PAM-DANs axons expressing GCaMP7f in the different MB compartments (GMR58E02-LexA > LexAop-GCaMP7f; A, anterior; P, posterior; scale bar=20µm). (**D**) Averaged ΔF/F calcium responses to vinegar-odor stimulation (*purple*) in the β2 compartment in fed and 24h starved control flies (->AdoRi) or in flies expressing *AdoR*-RNAi in PNG (*blue*; PNG>AdoRi) and CG (*orange*; CG>AdoRi), respectively. (**E**) Scatter plot of the maximal peak responses to vinegar-odor in the different groups (N_->AdoRi_=10/11; N_PNG>AdoRi_=8/7; N_CG>AdoRi_=8/7; 2-way ANOVA p(genotype)=0.12, p(state)=0.11, p(interaction)=0.0005). (**F**) Averaged ΔF/F responses to vinegar-odor stimulation (*purple*) in the γ5 compartment. (**G**) Scatter plot of the maximal peak responses to vinegar-odor in the different groups (2-way ANOVA p(genotype)=0.14, p(state)=0.31, p(interaction)=0.07). (**H**) Averaged ΔF/F responses to vinegar-odor stimulation (*purple*) in the γ4 compartment. (**I**) Scatter plot of the maximal peak responses to vinegar-odor in the different groups (2-way ANOVA p(genotype)=0.0349, p(state)=0.0362, p(interaction)=0.0018). (**J**) Averaged ΔF/F responses to vinegar-odor stimulation (*purple*) in the γ3 compartment. (**K**) Scatter plot of the maximal peak responses to vinegar-odor in the different groups (2-way ANOVA p(genotype)=0.0294, p(state)=0.0108, p(interaction)=0.25). Post hoc pair-wise comparisons are indicated as follow: ns, non-significant; *, p<0.05; **, p<0.01; *** p<0.001.

Taken together, these data show that adenosine signaling and its influence on glia modulates neuronal activity of DANs in a feeding state-dependent manner. Consistent with our behavioral observations, PNG and CG act in opposite ways to modulate DAN responses to vinegar-odor and in turn, conceivably contribute to the regulation of feeding and food-odor tracking behavior.

## Discussion

Physiological need induces a suite of behaviors such as foraging and feeding in food-deprived animals. While the role of neurons in regulating the expression of such behaviors has been amply documented, the function of the other large population of cell types in the brain, the glia, remains incompletely understood. In the present study, we show that adenosine and its receptor regulate the expression of feeding-related behaviors through different subtypes of glia cells. Importantly, these distinct subtypes of glial cells found in the fly CNS read this adenosine signal differently. On one hand, AdoR-mediated signaling in PNG, one of the two constituents of the fly HBB, and ALG specifically promotes feeding and foraging behavior, respectively, by increasing intracellular calcium levels (Figure 7B). On the other hand, adenosine signaling suppresses food-odor tracking and feeding by reducing calcium levels in CG upon feeding (Figure 7A). The opposite roles of PNG and CG are paralleled by opposite effects on DAN responses to vinegar-odor, thereby contributing to the expression of the appropriate behavior depending on the fly’s metabolic state.

**Figure 7:**
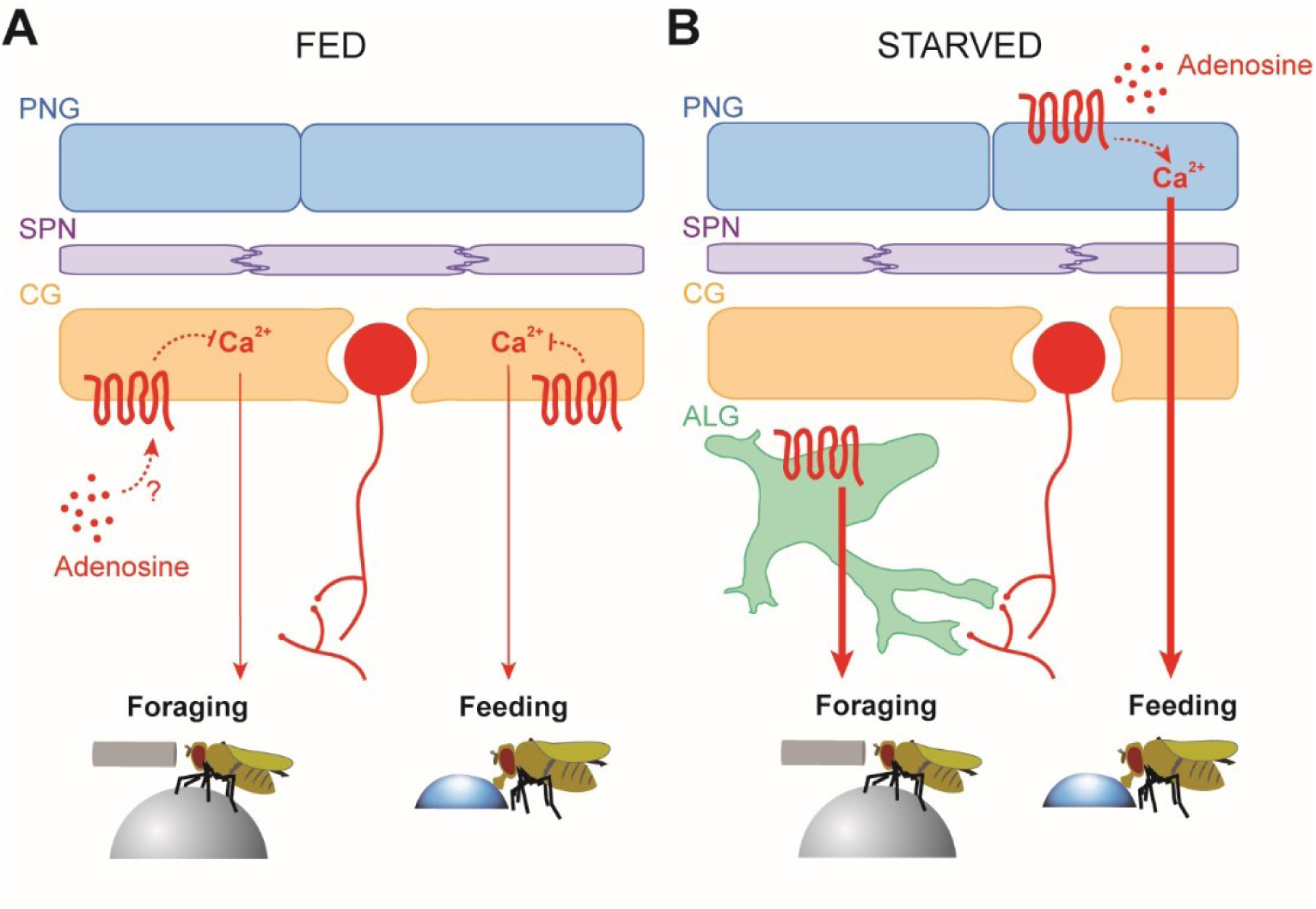
Summary model for the modulation of metabolic state-dependent behavior by adenosine signaling in glia. (**A**) Adenosine signaling prevents calcium increase in CG, refraining fed flies from tracking food-odor and feeding. (**B**) In starved flies, adenosine signaling in PNG promotes feeding behavior by increasing intracellular calcium concentration, while promoting food-odor tracking via ALG.

As one of the two constituents of the fly HBB, PNG has been shown to be crucial for the transfer of nutrients to fuel the nervous system, a role conserved in the vertebrate BBB^26,70^. Due to its anatomical location as the interface between the nutrient-containing hemolymph and the nutrient-consuming CNS, the HBB and glia in general have been proposed to function as a nutritional sensor capable of adapting the CNS to food shortages^27,29,30,71^. In addition, calcium transients have been recently reported in the fly PNG and seem to be important in contributing to the maintenance of neuronal excitability, suggesting that PNG could therefore influence behavior^53^. Our results provide support for this claim. We show that cytoplasmic calcium levels are increased in the PNG of starved flies in a way that depends on the presence of sugar in the extracellular saline solution. Calcium increases in PNG recorded in sugar-free solution was indeed abolished when trehalose and glucose, the main circulating types of sugar in the fly hemolymph, were added back to the saline, suggesting that circulating sugar maintains low calcium activity in the PNG of fed flies.

In addition to PNG, CG also plays an important role in nutrient storage in the CNS and in nutrient transfer to neurons to support their high demand in energy during the formation of long-term memory, for example^25,54,72–75^. This function is conserved in vertebrates and might resemble the classical, albeit still controversial, astrocyte-neuron lactate shuttle^27,76^. Furthermore, CG regulate neuronal excitability through its calcium activity^16,17^. We show here that this intracellular calcium level is modulated by the animal’s feeding state. Interestingly, however, the presence of trehalose and glucose does not seem to influence CG calcium levels, even though CG express glucose transporters^25,54^, suggesting that CG do not directly sense changes in hemolymph sugar concentration that occur during food deprivation. CG must therefore detect the metabolic state of the fly by other mechanisms, for example via insulin signaling^54^. CG has also been shown to detect its intracellular energetic state in response to starvation^73^. Our data show that adenosine signaling is necessary to maintain low calcium levels in the cytoplasm of CG in fed flies. Without AdoR, calcium levels in CG are increased and fed flies behave similarly to starved ones. Abnormally high calcium activity has previously been shown to elicit a dramatic increase in neuromuscular junction excitability recorded in fly larvae^16^. However, we rather observed an overall decrease in DANs response to vinegar-odor in the central brain. Several hypotheses could potentially explain this difference: (1) Calcium activity in CG might be heterogeneous throughout the CNS, as shown for PNG^53^, (2) The effect of calcium activity on neurons might be different in various regions of the CNS, and (3) The decreased excitability we observed in DANs might be indirect and a result of an alteration of activity in other parts of the neural circuit. Despite these remaining questions, our data indicate that CG cells detect the feeding state of the fly and help responds to starvation by transferring nutrients to support neuronal activity^73^ and by modulating neuronal activity and behavior. Interestingly, we also show in the present study that adenosine signaling is required in CG for the gain of weight induced by high fat, high sugar diet. Although the physiological mechanisms linking glia activity to energy storage remain to be elucidated, our data highlight the role of CNS glia in the regulation of systemic energetic metabolism, a function reminiscent of the role played by astrocytes in the mouse hypothalamus^28^.

Although our data primarily point toward a role for circulating adenosine in promoting feeding state-dependent behavior, we cannot exclude the participation of an independent, local production of extracellular adenosine in the brain of fed and/or starved flies. Indeed, we observed consequences of the knock-down of *AdoR* in glia intracellular calcium and in their effect on neuronal activity in both fed and starved flies. This suggests either a response of AdoR to a relatively low basal adenosine concentrations or a local increase in production/release of adenosine by neurons or glia. Interestingly, recent work from Themparambil and colleagues^77^ showed that active neurons release ATP that, after conversion into adenosine, signals to astrocytes to promote glucose uptake and subsequently release of lactate to support neuronal activity. This further highlights the role of adenosine signaling in the function of the neurovascular unit that fuels neurons in response to their energy demand^78^. We can therefore speculate that a local, activity-dependent release of ATP/adenosine may occur in the fly brain and signal through AdoR independently of systemic, circulating adenosine. Although tempting, this hypothesis remains to be demonstrated by further work.

Aside from its systemic and neuromodulatory roles, adenosine has been reported to act as a gliotransmitter in various model organism^31,79–83^. This could therefore potentially be an evolutionary well-conserved mechanism since the fly ALG has also been proposed to release ATP/adenosine in response to octopamine/tyramine -the fly functional analog of norepinephrine- to inhibit DAN activity and prevents chemotaxis in fly larvae^19^. We corroborate this role for adenosine signaling in DANs by showing that it inhibits feeding in starved flies, suggesting that adenosine might directly modulate neuronal circuits or through glia as an intermediate (Figure S2G-M). Further work using whole brain imaging combined with glia manipulation could provide additional insights to our understanding of glia function in the regulation of neural networks^69,84^.

### General remarks and conclusion

Being present and dispersed in the entire CNS, glial cells are in a good position to modulate neuronal activity on a large scale and have been shown to influence neuronal activity at the level of entire networks^8^. The ways glial cells can affect neurotransmission, and in turn behavior, are diverse and numerous. In mammals, a large variety of mechanisms used by astrocytes to modulate neuronal excitability and synaptic transmission have been described^6^. Interestingly, most of them found their equivalent in the fly’s glia, in ALG or in other cell types. This includes the release of signaling molecules^19,21,23^, the buffering of extra-cellular potassium^16,17^, as well as the transport and recycling of neurotransmitters^85,86^. Conversely to what it has been previously observed for the CG^16^ and the PNG^53^, we report a decrease in neuronal excitability in DANs correlated with an increase in calcium levels in those cell types. A decrease in odor-response has been shown following ALG activation in the antennal lobe^87^. This suggest that glia-mediated neuromodulation is more heterogeneous than previously shown in other flies studies and in line with current knowledge of astrocytes functions in mammals and other organisms^6,8^. Although we did not address those mechanisms in the present study, we showed that the alteration of calcium levels in two distinct sub-type of glial cells due to impaired adenosine signaling strongly affects the response of DANs to an appetitive odor.

In conclusion, in the present study, we identify adenosine signaling in glia as a key modulator that helps an animal adapt its behavior according to its internal metabolic state. The data presented here support the idea that purinergic signaling can provide information to the brain about decreasing internal nutrient availability in order to prioritize behaviors that will restore the energy balance of the organism. In addition, we provide the first evidences for the presence of purinergic signaling in fly glia, including in glia subtypes that comprise HBB which forms the interface between the nutrient-containing hemolymph and the CNS. Importantly, this function is differentially and specifically distributed among different glial cell types. This further highlights the role of the diversity and specialization of the fly glia subpopulations in the modulation of behavior, that all together recapitulate the glia functions^6^. Together, we suggest that purinergic signaling in glia and the BBB is a functionally conserved mechanism that is used by animals to sense their internal energetic state and adjust their behavior accordingly.

## Author contributions

J.-F.D.B. and I.C.G.K. conceptualized the study and designed the experiments. J.-F.D.B. carried out and analyzed the behavioral analysis with help from J.P. and T.K. J.-F.D.B. carried out and analyzed calcium imaging experiments. C.C. and R.A. designed, performed and analyzed LC-MS/MS analysis. Y.X. and C.G-C. designed, performed and analyzed TAG analysis. J.-F.D.B. and I.C.G.K. wrote the manuscript.

## Acknowledgments

We thank Heidi Miller-Mommerskamp and Natalie Lindenberg for help in general fly husbandry. We also thank Zeynep Pınar and Julia von Poblotzki for help with behavior experiments. We are grateful to Karen Y. Cheng, Annika Cichy and Aurélie Muria for their thoughtful comments on the manuscript.

We thank the German Research Foundation for funding this project (GR4310/11-1 to I.C.G.K.) and for funding essential research equipment (INST 217/1135-1 to I.C.G.K.).

## Supplemental information

Figures S1-S5

## Materials and Methods

### Flies strains and maintenance

Flies were raised on standard cornmeal food under 12:12 hours light/dark cycle on standard cornmeal food at 25°C and 60% humidity. For starvation experiments, flies were transferred to a starvation vial containing a wet tissue paper as water source 24hours prior experiments. For optogenetic experiments, adult flies were collected after hatching and kept on all trans-retinal supplemented food (1:250) under blue light only conditions. All experiments were conducted on 5-8-day old female flies, unless stated elsewhere.

### Key resources table

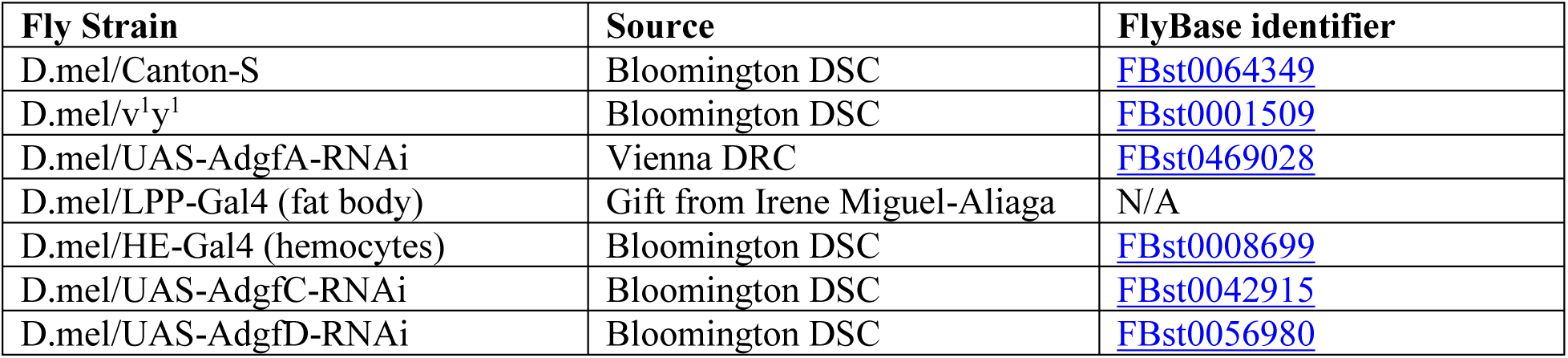
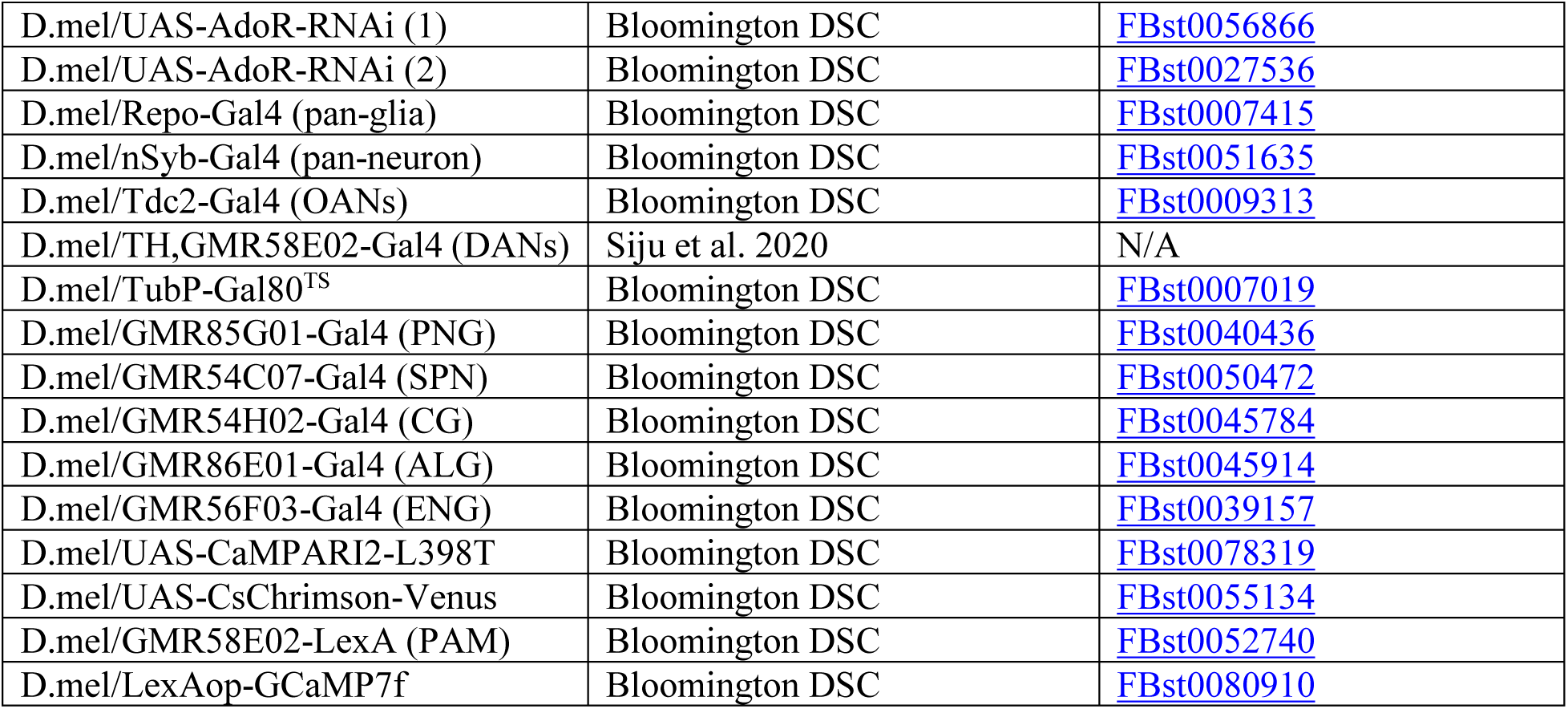

### FlyPAD feeding assay

FlyPAD feeding experiments were performed as previously described^46^. Single flies were transferred into the behavioral arena by mouth aspiration and left to feed for one hour on two 5µL drops of low-temperature melting agarose (1.5%) supplemented with 10% sucrose. Experiments were conducted in a climate chamber where a temperature of 25°C and 60% humidity were maintained. For optogenetic genetic experiments, we used the closed-loop stimulation paradigm described by Moreira et al.^62^. In this system, CsChrimson was activated by red light (625nm; 30µW/mm^2^) for a continuous period of 10 seconds when the fly’s proboscis contacted the drop of sucrose. Capacitance signal analysis was conducted using a custom-made MATLAB script provided by Pavel Itskov. The structure of feeding behavior was analyzed following the same parameters as described before^46^ (Figure 1B). The cumulated and total number of sips, number of activity bouts and number of feeding bursts were summed for the two electrodes. The sip durations, inter-sips intervals, durations of activity bouts and durations of feeding bursts were averaged for the two electrodes.

### Spherical treadmill behavioral assay

Single fly experiments on the spherical treadmill were performed using the same device and paradigm as previously described^50^. Briefly, the experimental fly was anesthetized on ice and tethered by gluing a tungsten pin on their thorax using UV-cured clue. Another drop of glue was used to glue the posterior part of the head to anterior part of the thorax and the wings were clipped. The fly was then immediately transferred onto the treadmill and was given 3 minutes of acclimatization before experiment initialization. Each trial consisted in pre-stimulation (20s), stimulation with balsamic vinegar odor (4ppm; 12s) and post-stimulation periods (30s; Figure 2I). The 10 consecutive recorded trials were separated by semi-randomized inter-trial periods of 60 ± 2-20 seconds. Data acquisition an analysis were performed using custom-written Python 2.7 scripts, as previously described^50^. For analysis, we measured the average running speed of the fly on the spherical treadmill during the 12s of odor stimulation.

### Adenosine content

#### Metabolite extraction

10 frozen flies per sample were transferred into Precellys® tubes prefilled with ceramic beads (hard tissue lysing kit, CK28-R) and 1mL of −20°C MeOH:H_2_O (4:1, v/v) was added to each sample. The homogenization was performed with a Precellys® Evolution tissue homogenizer connected to a Cryolys® module. The temperature was maintained at 4°C while 4 homogenization cycles where performed at 7500 rpm for 20 s with 30 s break intervals. Next, homogenates were transferred into Eppendorf tubes, incubated for 15 min at 4°C and 950 rpm and finally centrifuged for 15 min at 15,500 rpm and 4°C. The recovered supernatant was dried under a gentle nitrogen flow. Each sample was reconstituted in 50 µL AcN/H_2_O (9:1, v/v), briefly sonicated and centrifuged before LC separation.

#### LC-MS/MS analysis

Analysis of adenosine was performed as previously described with minor modifications^88,89^. Briefly, a Vanquish Flex UHPLC system (Thermo Fisher Scientific, Germering, Germany) was equipped with a fitted iHILIC-(P) Classic (2.1 x 100 mm, 5 µm) column and pre-column (20 x 2.1 mm, 5 μm, 200 Å; both Dichrom, Haltern am See, Germany). The mobile phases were 15 mM ammonium acetate in H_2_O at pH = 9.4 (A) and 100% acetonitrile (B). Separation was achieved with the following gradient: 0 to 12 min ramp from 90% B to 26% B, 12 to 14 min 26% B, 14 to 17 minutes ramp to 10% B, 17 to19 min 10% B, 19 to 19.1 min ramp back to 90% B and equilibrate until min 27. 5 µL of each sample were injected onto the column twice (spiked/not spiked with an isotopically labeled adenosine standard (Eurisotop) and the LC separation was carried out at 40°C with a flow rate of 200 µL/min.

The LC system was coupled to a QTRAP 6500+ (Applied Biosystems, Darmstadt, Germany). The measurements were performed in positive mode with the following ESI Turbo V ion source parameters: curtain gas: 30 arbitrary units; temperature: 350°C; ion source gas 1:40 arbitrary units; ion source gas 2:65 arbitrary units; collision gas: medium; ion spray voltage: 5500 V. For the selected reaction monitoring (SRM) mode Q1 and Q3 were set to unit resolution and CE was 27 V. Data were acquired with Analyst (version 1.7.2; AB Sciex) and Skyline (3) was used to visualize results and manually integrate signals.

### TAG content

50 5-days old adult male flies were transferred to food vials containing either fly HFHS diet or normal fly food for 2 days, respectively. HFHS diet (∼20% fat, ∼12.5% sugar) consisted in: peanut butter 37%, coconut oil 1%, sucrose 7.5%, water 7.5%, standard fly food 45%, grape juice 2% and propionic acid 0.2% (v/v). To allow fly access to water and avoid them sticking to the HFHS diet during feeding, a portion of fly food was loaded on the top of 5ml agar (1% in water) in the vials. 5 male flies per replicate were collected and stored at −20 °C for fly body fat content measurement. Fly body fat content was determined by normalizing triglyceride to protein content of fly homogenates: triglyceride by the coupled-colorimetric assay (T7532, Pointe Scientific^90^) and protein by Bicinchoninic acid assay (23225, Thermofisher Scientific), as previously described^91,92^.

### *Ex vivo* calcium imaging

Fly brains were dissected in adult hemolymph-like saline (AHL; NaCl, 103mM; NaHCO_3_, 26mM; KCl, 3mM; CaCl_2_, 1.5mM; MgCl_2_, 4mM; NaH_2_PO_4_, 1mM; TES, 5mM; D-Glucose, 10mM; Trehalose, 10mM). In experiments simulating the hemolymph of starved flies, D-Glucose and Trehalose were replaced by 20mM D-ribose to preserve osmolarity. A single dissected brain was then mounted caudal side up between a microscope glass slide and a cover-slip using the same medium as for dissection and placed under an inverted Leica SP5 laser scanning confocal microscope. The photoconversion of CaMPARI2-L398T proteins expressed by glial cells was achieved by illuminating the sample for 30 seconds using the mercury lamp of the microscope and a 395/25nm filter through a 20x water immersion objective (NA 0.7). 512×512 pixel confocal stacks of approximately 20µm (CG) or 10µm (PNG) tick volume of tissue (Δz=1µm) were taken directly after photoconversion using the same 20x objective and a 4x digital zoom. The signal-to-noise ratio was improved by averaging 4 lines scans. An additional 405nm excitation laser was used in combination with the 488nm laser while recording the green channel to preserve CaMPARI2-L398T photoconversion^56^. Both CG and PNG were imaged on the dorso-caudal part of a randomly selected brain hemisphere, where the calyx of the MB is clearly identifiable. Data were analyzed using ImageJ (NIH) by manual drawing ROIs on z-maximal projections of the confocal stacks. The ratio between the florescence intensities for the red and green channels were averaged between the ROIs for a single fly and compared between the different experimental conditions.

### *In vivo* 2-photon calcium imaging

For *in vivo* imaging, flies were prepared as previously described^66^. In this preparation, the fly’s body was restrained in a truncated 1mL pipette tip, impairing its movements. After dissection of the head cuticle, fat body and tracheae, the exposed brain was covered by a drop of low-temperature melting agarose diluted in the recording AHL to prevent brain movements. Fed flies were recorded using standard AHL (NaCl, 103mM; NaHCO_3_, 26mM; KCl, 3mM; CaCl_2_, 1.5mM; MgCl_2_, 4mM; NaH_2_PO_4_, 1mM; TES, 5mM; D-Glucose, 10mM; Trehalose, 10mM), while starved flies were recorded using a solution depleted in D-Glucose and Trehalose (NaCl, 103mM; NaHCO_3_, 26mM; KCl, 3mM; CaCl_2_, 1.5mM; MgCl_2_, 4mM; NaH_2_PO_4_; 1mM, TES, 5mM; D-ribose, 10mM).

GCaMP7f fluorescence signal was recorded on a Rapp OptoElectronic customized Sutter Instrument MOM 2-photon microscope equipped with a 25x Nikon water immersion objective (NA 1.10), 8kHz RGG resonant scanner, a fast z-piezo and a Hammamatsu gateable PMT. Excitation light was provided by a 920mm Toptica photonics laser. Stacks comprising the axons terminals of the PAM neurons innervating the horizontal lobe of the mushroom body were imaged at a frequency of 2.31Hz (512×512 pixels; digital zoom 4x; 13 slices; Δz=4µm).

Vinegar-odor stimulation (1% in water) was delivered through a Syntech stimulus controller CS-55. During the entire duration of the experiment, flies were exposed to a continuous air flow (1mL/min). Flies were stimulated for 12 seconds after a baseline recording of 30 seconds, to match with our spherical treadmill paradigm.

For analysis, ROIs delimiting the different lobes of the mushroom body (β2, γ3, 4 and 5) were manually drown using ImageJ (NIH) on a maximum z-projection of the imaged stacks, for each trial. Raw traces were then analyzed using custom Python3.0 scripts. For β2 and γ4 compartments, relative GCaMP fluorescence intensities (ΔF/F_0_) were computed using the averaged fluorescence intensity for the 5 frames preceding the stimulus presentation (F_0_). The vinegar-response peak was defined as the maximal ΔF/F_0_ value measured during the stimulus phase. Since the γ3 and 5 compartments did not show a clear baseline but rather an oscillatory pattern of activity, we calculated the basal fluorescence (F_0_) by averaging the fluorescence intensity over the entire trial recording. The vinegar response and spontaneous peaks were defined as the difference between the maximal ΔF/F_0_ value measured during the stimulus phase and the minimal ΔF/F_0_ value measured directly before stimulation. The amplitude of the spontaneous peaks was averaged for each trial.

### Statistical analysis

Results are expressed as the mean ± SEM or as a scatter plot displaying the individual data point as well as their mean ± SEM. Data significance was tested by Mann-Whitney U-test, Welch’s t-test, one-way ANOVA, two-way ANOVA, two-way RM ANOVA or three-way ANOVA, depending on the experiment design. Details related to sample sizes, statistical tests and post hoc pairwise comparisons used for statistical analysis are summarized in the figure legends. The reported sample sizes are the number of flies used for the given experiment, except for the measurements of adenosine and TAG content where the number of flies per replicate is stated. Statistical analysis was performed using GraphPad Prism 10. The significance threshold was set as p-value>0.05.

## Supplemental Information

**Figure S1:**
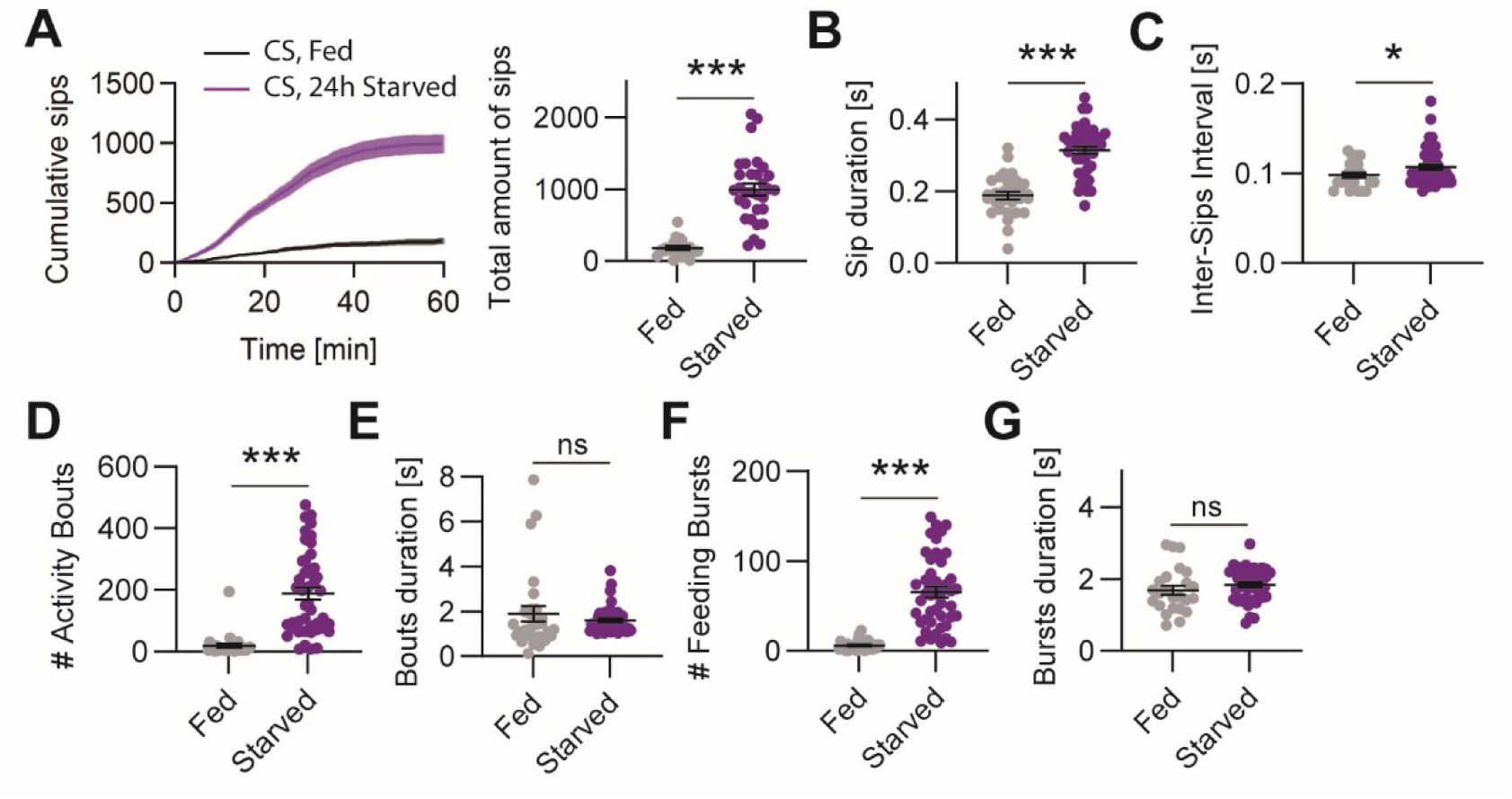
Feeding behavior in fed and 24h starved wild-type flies (related to Figure 1) (**A**) Cumulative number of sips (mean ± sem) and scatter plot of the total number sips on 10% sucrose drops in fed and 24h starved CS flies in one hour (N=29/45; Welch’s t-test p<0.0001). (**B**) Averaged sip duration (Welch’s t-test p<0.0001). (**C**) Averaged time interval between sips (Welch’s t-test p=0.0459). (**D**) Total number of activity bouts (Welch’s t-test p<0.0001). (**E**) Averaged duration of activity bouts (Welch’s t-test p=0.40). (**F**) Total number of feeding bursts (Welch’s t-test p<0.0001). (**G**) Averaged duration of feeding bursts (Welch’s t-test p=0.29).

**Figure S2:**
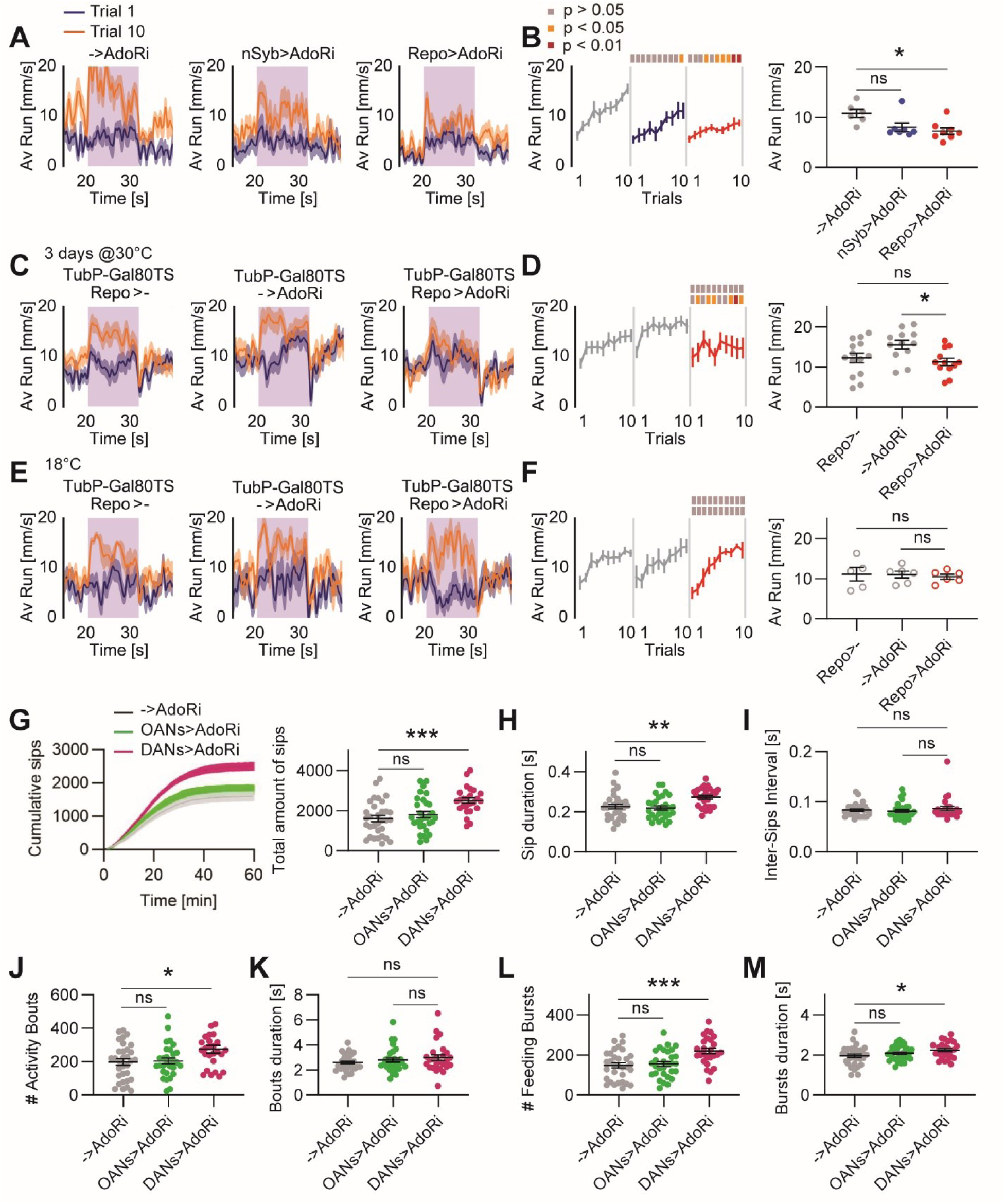
Adenosine signaling in dopaminergic neurons inhibits feeding behavior (related to Figure 2) (**A**) Averaged forward running speed (± sem) of 24h starved control flies (->AdoRi) or upon knock-down of AdoR in neurons (nSyb>AdoRi) and glia (Repo>AdoRi) using an alternative RNAi line, respectively. (**B**) Running speed during the stimulus phase over 10 successive trials (mean ± sem; N=6/7/8; 2-way RM ANOVA p(groups)=0.0147; p(trials)<0.0001; p(interaction)=0.0830). (**C-F**) Restriction of the AdoR knock-down in adult flies using the temperature-sensitive TARGET system. (**C**) Averaged forward running speed (± sem) of 24h starved control flies (TubP-Gal80^TS^, Repo>- and TubP-Gal80^TS^, ->AdoRi) or upon knock-down of AdoR in glia (TubP-Gal80^TS^, Repo>AdoRi) in flies raised at 18°C to prevent RNAi expression. Flies were kept for 2 days at 30°C before experiment to enable RNAi expression. (**D**) Running speed during the stimulus phase over 10 successive trials (mean ± sem; N=14/12/12; 2-way RM ANOVA p(groups)=0.0275; p(trials)<0.0001; p(interaction)=0.54). (**E**) Averaged forward running speed (± sem) of 24h starved control flies (TubP-Gal80^TS^, Repo>- and TubP-Gal80^TS^, ->AdoRi) or upon knock-down of AdoR in glia (TubP-Gal80^TS^, Repo>AdoRi) in flies raised and kept at 18°C to prevent RNAi expression. (**F**) Running speed during the stimulus phase over 10 successive trials (mean ± sem; N=6/5/6; 2-way RM ANOVA p(groups)=0.90; p(trials)<0.0001; p(interaction)=0.13). (**G**) Cumulative number of sips (mean ± sem) and scatter plot of the total number sips on 10% sucrose drops in 24h starved control flies (->AdoRi) or upon knock-down of AdoR in octopaminergic (OANs>AdoRi, using Tdc2-Gal4) and dopaminergic neurons (DANs>AdoRi, using TH,GMR58E02-Gal4; N=32/30/25; one-way ANOVA p=0.0002). (**H**) Averaged sip duration (one-way ANOVA p=0.0002). (**I**) Averaged time interval between sips (one-way ANOVA p=0.56). (**J**) Total number of activity bouts (one-way ANOVA p=0.0179). (**K**) Averaged duration of activity bouts (one-way ANOVA p=0.28). (**L**) Total number of feeding bursts (one-way ANOVA p=0.0004). (**M**) Averaged duration of feeding bursts (one-way ANOVA p=0.0437). For food-odor tracking behavior experiments, Sidak’s post hoc trial-to-trial comparisons are depicted on the top of the graphs as color-coded boxes (grey p>0.05, orange p<0.05 and red p<0.01). The scatter plots represent the main group effect of the ANOVA. Post hoc pair-wise comparisons are indicated as follow: ns, non-significant; *, p<0.05; **, p<0.01; *** p<0.001.

**Figure S3:**
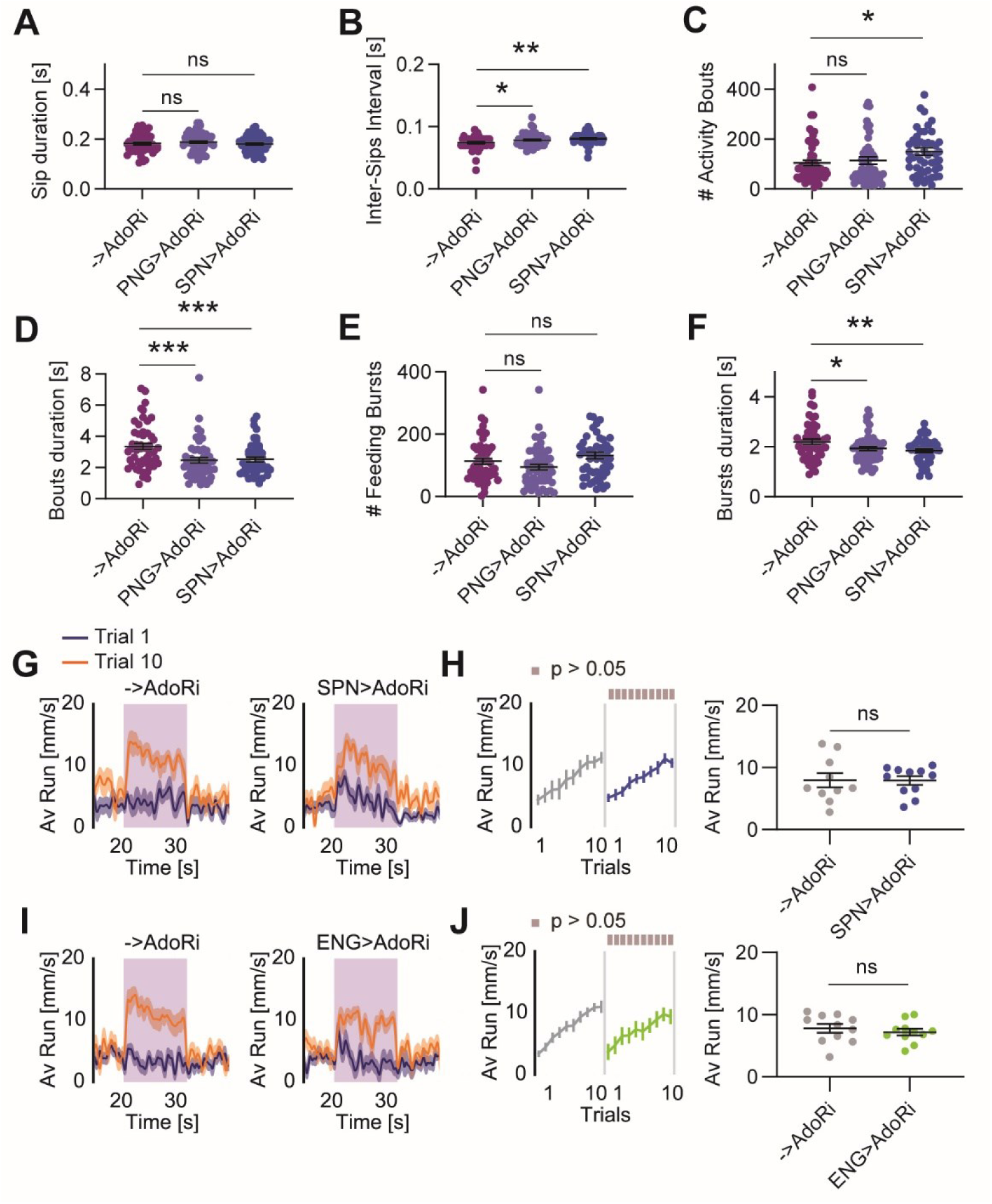
Adenosine signaling in subperineurial and ensheating glia is not necessary for food-odor tracking behavior (related to Figure 3) (**A**) Averaged sip duration in 24h starved control flies (->AdoRi) or flies deficient for AdoR in PNG (PNG>AdoRi) and SPN (SPN>AdoRi), respectively (N=49/51/48; one-way ANOVA p=0.60). (**B**) Averaged time interval between sips (one-way ANOVA p=0.0029). (**C**) Total number of activity bouts (one-way ANOVA p=0.0506). (**D**) Averaged duration of activity bouts (one-way ANOVA p=0.0004). (**E**) Total number of feeding bursts (one-way ANOVA p=0.0342). (**F**) Averaged duration of feeding bursts (one-way ANOVA p=0.0111). (**G**) Averaged forward running speed (± sem) of 24h starved control flies (->AdoRi) or upon knock-down of AdoR in SPN (SPN>AdoRi) (**H**) Running speed during the stimulus phase over 10 successive trials (mean ± sem; N=10/11; 2-way RM ANOVA p(groups)=0.98; p(trials)<0.0001; p(interaction)=0.85). (**I**) Averaged forward running speed (± sem) of 24h starved control flies (->AdoRi) or upon knock-down of AdoR in ENG (ENG>AdoRi). (**J**) Running speed during the stimulus phase over 10 successive trials (mean ± sem; N=10/11; 2-way RM ANOVA p(groups)=0.47; p(trials)<0.0001; p(interaction)=0.77). For food-odor tracking behavior experiments, Sidak’s post hoc trial-to-trial comparisons are depicted on the top of the graphs as color-coded boxes (grey p>0.05, orange p<0.05 and red p<0.01). The scatter plots represent the main group effect of the ANOVA. Post hoc pair-wise comparisons are indicated as follow: ns, non-significant; *, p<0.05; **, p<0.01; *** p<0.001.

**Figure S4:**
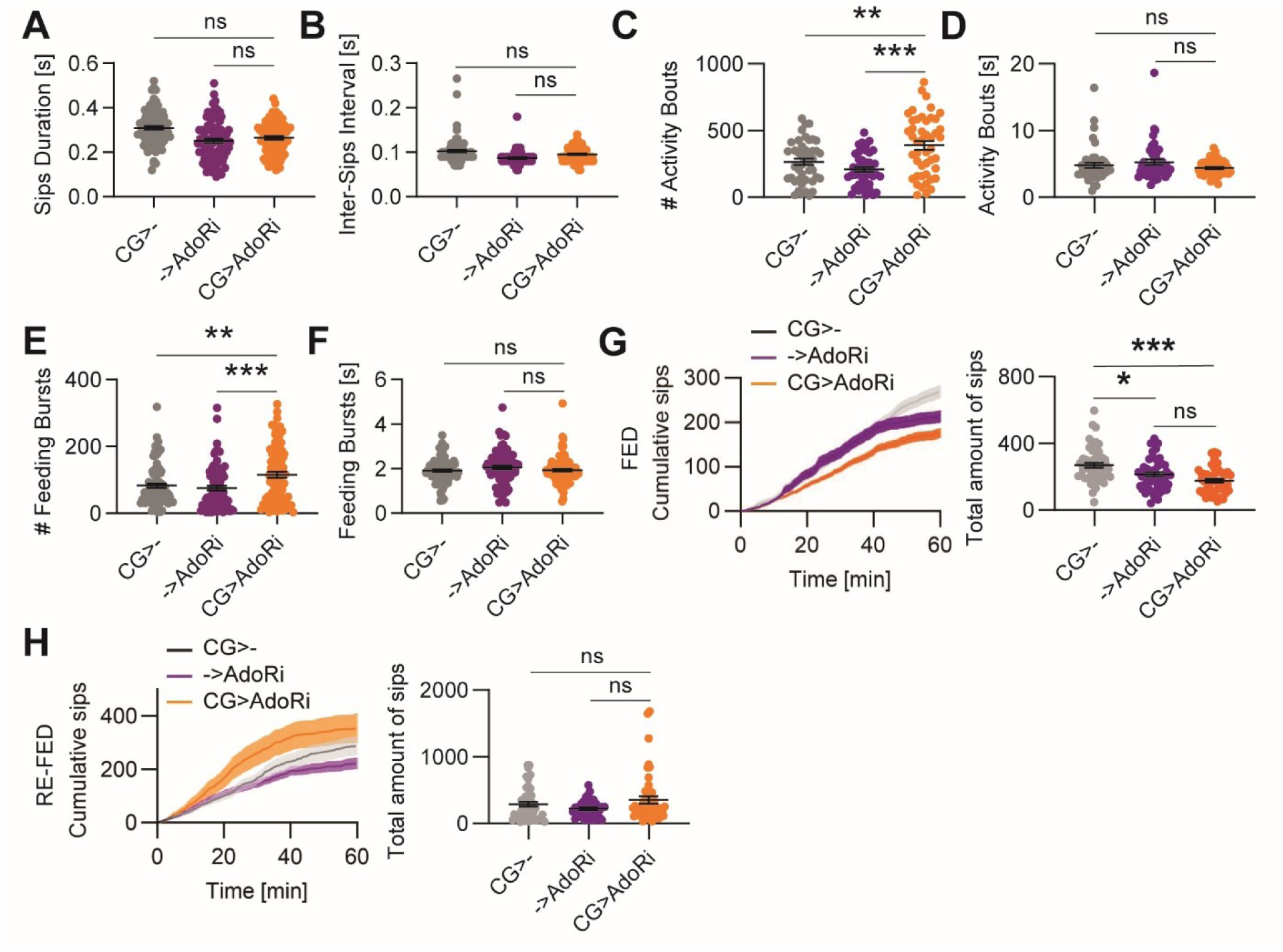
Adenosine signaling in cortex glia inhibits feeding and food-odor tracking behavior (Related to Figure 4) (**A**) Averaged sip duration in 24h starved control flies (CG>- and ->AdoRi) or flies deficient for AdoR in CG (CG>AdoRi; N=45/44/44; one-way ANOVA p<0.0001). (**B**) Averaged time interval between sips (one-way ANOVA p<0.0001). (**C**) Total number of activity bouts (one-way ANOVA p<0.0001). (**D**) Averaged duration of activity bouts (one-way ANOVA p=0.22). (**E**) Total number of feeding bursts (one-way ANOVA p=0.0003). (**F**) Averaged duration of feeding bursts (one-way ANOVA p=0.29). (**G**) Cumulative number of sips (mean ± sem) and scatter plot of the total number sips on drops of 10% sucrose in fed control flies (CG>- and ->AdoRi) or upon knock-down of AdoR in CG (CG>AdoRi; N=53/48/48; one-way ANOVA p<0.0001). (**H**) Cumulative number of sips (mean ± sem) and scatter plot of the total number sips on 10% sucrose drops in 24h starved control flies (CG>- and ->AdoRi) or AdoR-deficient flies in CG (CG>AdoRi) re-fed for 40 minutes on standard fly food (N=46/36/47; Brown-Forsythe one-way ANOVA p=0.0841).

**Figure S5:**
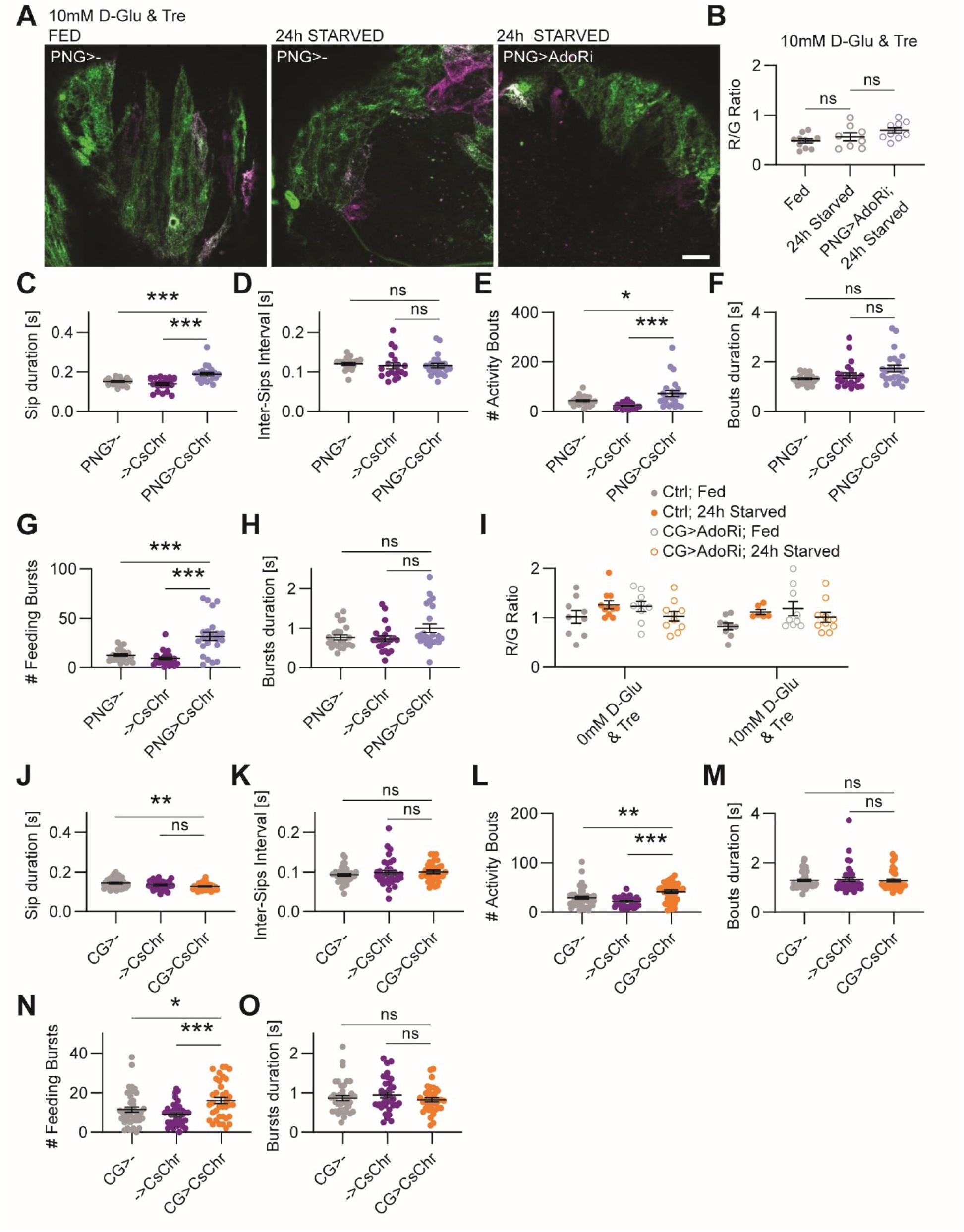
Starvation and adenosine signaling modulate intra-cellular calcium levels in perineurial and cortex glia (related to Figure 5) (**A**) Example images of CaMPARI2-L398T-expressing PNG cells in dorso-caudal brain explants from fed and 24h starved control flies (PNG>-) and upon knock-down of AdoR (PNG>AdoRi), in standard AHL (10mM D-Glu and 10mM Tre), after photoconversion (scale bar=20µm). (**B**) Scatter plot of photoconversion ratios in fed and 24h starved control flies (N=11/8), as well as 24h starved PNG>AdoRi flies (N=10; 2-way ANOVA p(AHL)=0.0076; p(groups)=0.06; p(interaction)=0.06). (**C**) Averaged sip duration in fed flies upon closed-loop optogenetic activation of PNG (N=23/22/23; one-way ANOVA p<0.0001). (**D**) Averaged time interval between sips (one-way ANOVA p=0.80). (**E**) Total number of activity bouts (one-way ANOVA p=0.0002). (**F**) Averaged duration of activity bouts (one-way ANOVA p=0.0152). (**G**) Total number of feeding bursts (one-way ANOVA p<0.0001). (**H**) Averaged duration of feeding bursts (one-way ANOVA p=0.06). (**I**). Scatter plot of photoconversion ratios in fed and 24h starved control and CG>AdoRi flies imaged in standard (N_fed_=9/11; N_starved_=9/10) or sugar-free AHL (N_fed_=9/8; N_starved_=9/10; 3-way ANOVA p(state)=0.43; p(genotype)=0.59; p(AHL)=0.18; p(state x genotype)=0.0022; p(state x AHL)=0.35; p(genotype x AHL)=0.79; p(state x genotype x AHL)=0.95). Data from standard and sugar-free AHL are pooled for display in Figure 5F. (**J**) Averaged sip duration in fed flies upon closed-loop optogenetic activation of CG (N=44/41/37; one-way ANOVA p=0.0037). (**K**) Averaged time interval between sips (one-way ANOVA p=0.47). (**L**) Total number of activity bouts (one-way ANOVA p<0.0001). (**M**) Averaged duration of activity bouts (one-way ANOVA p=0.85). (**N**) Total number of feeding bursts (one-way ANOVA p=0.0012). (**O**) Averaged duration of feeding bursts (one-way ANOVA p=0.41). Post hoc pair-wise comparisons are indicated as follow: ns, non-significant; *, p<0.05; **, p<0.01; *** p<0.001.

## References

1. Rolls, E.T. (2006). Brain mechanisms underlying flavour and appetite. Philos. Trans. R. Soc. B Biol. Sci. 361, 1123–1136. 10.1098/rstb.2006.1852

2. Itskov, P.M., and Ribeiro, C. (2013). The Dilemmas of the Gourmet Fly: The Molecular and Neuronal Mechanisms of Feeding and Nutrient Decision Making in Drosophila. Front. Neurosci. 7. 10.3389/fnins.2013.00012.

3. Cichy, A. (2022). How the body rules the nose. Neuroforum 28, 151–158. 10.1515/nf-2022-0003.

4. Sachse, S., and Beshel, J. (2016). The good, the bad, and the hungry: how the central brain codes odor valence to facilitate food approach in Drosophila. Curr. Opin. Neurobiol. 40, 53–58. 10.1016/j.conb.2016.06.012.

5. Sayin, S., Boehm, A.C., Kobler, J.M., De Backer, J.-F., and Grunwald Kadow, I.C. (2018). Internal State Dependent Odor Processing and Perception—The Role of Neuromodulation in the Fly Olfactory System. Front. Cell. Neurosci. 12, 11. 10.3389/fncel.2018.00011.

6. Nagai, J., Yu, X., Papouin, T., Cheong, E., Freeman, M.R., Monk, K.R., Hastings, M.H., Haydon, P.G., Rowitch, D., Shaham, S., et al. (2021). Behaviorally consequential astrocytic regulation of neural circuits. Neuron 109, 576–596. 10.1016/j.neuron.2020.12.008.

7. Bazargani, N., and Attwell, D. (2016). Astrocyte calcium signaling: the third wave. Nat. Neurosci. 19, 182–189. 10.1038/nn.4201.

8. Araque, A., Carmignoto, G., Haydon, P.G., Oliet, S.H.R., Robitaille, R., and Volterra, A. (2014). Gliotransmitters travel in time and space. Neuron 81, 728–739. 10.1016/j.neuron.2014.02.007.

9. Yang, L., Qi, Y., and Yang, Y. (2015). Astrocytes Control Food Intake by Inhibiting AGRP Neuron Activity via Adenosine A1 Receptors. Cell Rep. 11, 798–807. 10.1016/j.celrep.2015.04.002.

10. Chen, N., Sugihara, H., Kim, J., Fu, Z., Barak, B., Sur, M., Feng, G., and Han, W. (2016). Direct modulation of GFAP-expressing glia in the arcuate nucleus bi-directionally regulates feeding. eLife 5, e18716. 10.7554/eLife.18716.

11. García-Cáceres, C., Balland, E., Prevot, V., Luquet, S., Woods, S.C., Koch, M., Horvath, T.L., Yi, C.-X., Chowen, J.A., Verkhratsky, A., et al. (2019). Role of astrocytes, microglia, and tanycytes in brain control of systemic metabolism. Nat. Neurosci. 22, 7–14. 10.1038/s41593-018-0286-y.

12. Herrera Moro Chao, D., Kirchner, M.K., Pham, C., Foppen, E., Denis, R.G.P., Castel, J., Morel, C., Montalban, E., Hassouna, R., Bui, L.-C., et al. (2022). Hypothalamic astrocytes control systemic glucose metabolism and energy balance. Cell Metab. 34, 1532–1547.e6. 10.1016/j.cmet.2022.09.002.

13. Zwarts, L., Van Eijs, F., and Callaerts, P. (2015). Glia in Drosophila behavior. J. Comp. Physiol. A 201, 879–893. 10.1007/s00359-014-0952-9.

14. Kremer, M.C., Jung, C., Batelli, S., Rubin, G.M., and Gaul, U. (2017). The glia of the adult *Drosophila* nervous system: Glia Anatomy in Adult Drosophila Nervous System. Glia 65, 606–638. 10.1002/glia.23115.

15. Yildirim, K., Petri, J., Kottmeier, R., and Klämbt, C. (2019). Drosophila glia: Few cell types and many conserved functions: YILDIRIM et al. Glia 67, 5–26. 10.1002/glia.23459.

16. Melom, J.E., and Littleton, J.T. (2013). Mutation of a NCKX Eliminates Glial Microdomain Calcium Oscillations and Enhances Seizure Susceptibility. J. Neurosci. 33, 1169–1178. 10.1523/JNEUROSCI.3920-12.2013.

17. Weiss, S., Melom, J.E., Ormerod, K.G., Zhang, Y.V., and Littleton, J.T. (2019). Glial Ca2+signaling links endocytosis to K+ buffering around neuronal somas to regulate excitability. eLife 8, e44186. 10.7554/eLife.44186.

18. Basu, R., Preat, T., and Plaçais, P.-Y. (2024). Glial metabolism versatility regulates mushroom body-driven behavioral output in Drosophila. Learn. Mem. Cold Spring Harb. N 31, a053823. 10.1101/lm.053823.123.

19. Ma, Z., Stork, T., Bergles, D.E., and Freeman, M.R. (2016). Neuromodulators signal through astrocytes to alter neural circuit activity and behaviour. Nature 539, 428–432. 10.1038/nature20145.

20. Blum, I.D., Keleş, M.F., Baz, E.-S., Han, E., Park, K., Luu, S., Issa, H., Brown, M., Ho, M.C.W., Tabuchi, M., et al. (2021). Astroglial Calcium Signaling Encodes Sleep Need in Drosophila. Curr. Biol. 31, 150–162.e7. 10.1016/j.cub.2020.10.012.

21. Park, A., Croset, V., Otto, N., Agarwal, D., Treiber, C.D., Meschi, E., Sims, D., and Waddell, S. (2022). Gliotransmission of D-serine promotes thirst-directed behaviors in Drosophila. Curr. Biol. 32, 3952–3970.e8. 10.1016/j.cub.2022.07.038.

22. Flores-Valle, A., and Seelig, J.D. (2022). Dynamics of sleep and feeding homeostasis in *Drosophila* glia and neurons. Preprint, 10.1101/2022.07.07.499175 10.1101/2022.07.07.499175.

23. Miyashita, T., Murakami, K., Kikuchi, E., Ofusa, K., Mikami, K., Endo, K., Miyaji, T., Moriyama, S., Konno, K., Muratani, H., et al. (2023). Glia transmit negative valence information during aversive learning in *Drosophila*. Science 382, eadf7429. 10.1126/science.adf7429.

24. O’Brown, N.M., Pfau, S.J., and Gu, C. (2018). Bridging barriers: a comparative look at the blood– brain barrier across organisms. Genes Dev. 32, 466–478. 10.1101/gad.309823.117.

25. Volkenhoff, A., Hirrlinger, J., Kappel, J.M., Klämbt, C., and Schirmeier, S. (2018). Live imaging using a FRET glucose sensor reveals glucose delivery to all cell types in the Drosophila brain. J. Insect Physiol. 106, 55–64. 10.1016/j.jinsphys.2017.07.010.

26. Volkenhoff, A., Weiler, A., Letzel, M., Stehling, M., Klämbt, C., and Schirmeier, S. (2015). Glial Glycolysis Is Essential for Neuronal Survival in Drosophila. Cell Metab. 22, 437–447. 10.1016/j.cmet.2015.07.006.

27. Rittschof, C.C., and Schirmeier, S. (2018). Insect models of central nervous system energy metabolism and its links to behavior. Glia 66, 1160–1175. 10.1002/glia.23235.

28. Murat, C.D.B., and García-Cáceres, C. (2021). Astrocyte Gliotransmission in the Regulation of Systemic Metabolism. Metabolites 11, 732. 10.3390/metabo11110732.

29. De Backer, J.-F., and Grunwald Kadow, I.C. (2022). A role for glia in cellular and systemic metabolism: insights from the fly. Curr. Opin. Insect Sci. 53, 100947. 10.1016/j.cois.2022.100947.

30. Contreras, E.G., and Sierralta, J. (2022). The Fly Blood-Brain Barrier Fights Against Nutritional Stress. Neurosci. Insights 17, 263310552211202. 10.1177/26331055221120252.

31. Lezmy, J. (2023). How astrocytic ATP shapes neuronal activity and brain circuits. Curr. Opin. Neurobiol. 79, 102685. 10.1016/j.conb.2023.102685.

32. Yegutkin, G.G. (2008). Nucleotide- and nucleoside-converting ectoenzymes: Important modulators of purinergic signalling cascade. Biochim. Biophys. Acta 1783, 673–694. 10.1016/j.bbamcr.2008.01.024.

33. Buck, L.T. (2004). Adenosine as a signal for ion channel arrest in anoxia-tolerant organisms. Comp. Biochem. Physiol. B Biochem. Mol. Biol. 139, 401–414. 10.1016/j.cbpc.2004.04.002.

34. Newby, A.C. (1984). Adenosine and the concept of ‘retaliatory metabolites.’ Trends Biochem. Sci. 9, 42–44. 10.1016/0968-0004(84)90176-2.

35. Haskó, G., Deitch, E.A., Szabó, C., Németh, Z.H., and Vizi, E.S. (2002). Adenosine: a potential mediator of immunosuppression in multiple organ failure. Curr. Opin. Pharmacol. 2, 440–444. 10.1016/s1471-4892(02)00172-8.

36. Davis, J.M., Zhao, Z., Stock, H.S., Mehl, K.A., Buggy, J., and Hand, G.A. (2003). Central nervous system effects of caffeine and adenosine on fatigue. Am. J. Physiol. Regul. Integr. Comp. Physiol. 284, R399–404. 10.1152/ajpregu.00386.2002.

37. Antonioli, L., Csóka, B., Fornai, M., Colucci, R., Kókai, E., Blandizzi, C., and Haskó, G. (2014). Adenosine and inflammation: what’s new on the horizon? Drug Discov. Today 19, 1051–1068. 10.1016/j.drudis.2014.02.010.

38. Koupenova, M., and Ravid, K. (2013). Adenosine, Adenosine Receptors and Their Role in Glucose Homeostasis and Lipid Metabolism. J. Cell. Physiol. 228, 1703–1712. 10.1002/jcp.24352.

39. Verkhratsky, A., and Burnstock, G. (2014). Biology of purinergic signalling: Its ancient evolutionary roots, its omnipresence and its multiple functional significance. BioEssays 36, 697–705. 10.1002/bies.201400024.

40. Zuberova, M., Fenckova, M., Simek, P., Janeckova, L., and Dolezal, T. (2010). Increased extracellular adenosine in Drosophila that are deficient in adenosine deaminase activates a release of energy stores leading to wasting and death. Dis. Model. Mech. 3, 773–784. 10.1242/dmm.005389.

41. Bajgar, A., Kucerova, K., Jonatova, L., Tomcala, A., Schneedorferova, I., Okrouhlik, J., and Dolezal, T. (2015). Extracellular adenosine mediates a systemic metabolic switch during immune response. PLoS Biol. 13, e1002135. 10.1371/journal.pbio.1002135.

42. Bajgar, A., and Dolezal, T. (2018). Extracellular adenosine modulates host-pathogen interactions through regulation of systemic metabolism during immune response in Drosophila. PLOS Pathog. 14, e1007022. 10.1371/journal.ppat.1007022.

43. Zurovec, M., Dolezal, T., Gazi, M., Pavlova, E., and Bryant, P.J. (2002). Adenosine deaminase-related growth factors stimulate cell proliferation in *Drosophila* by depleting extracellular adenosine. Proc. Natl. Acad. Sci. 99, 4403–4408. 10.1073/pnas.062059699.

44. Dolezal, T., Dolezelova, E., Zurovec, M., and Bryant, P.J. (2005). A Role for Adenosine Deaminase in Drosophila Larval Development. PLoS Biol. 3, e201. 10.1371/journal.pbio.0030201.

45. Novakova, M., and Dolezal, T. (2011). Expression of Drosophila Adenosine Deaminase in Immune Cells during Inflammatory Response. PLoS ONE 6, e17741. 10.1371/journal.pone.0017741.

46. Itskov, P.M., Moreira, J.-M., Vinnik, E., Lopes, G., Safarik, S., Dickinson, M.H., and Ribeiro, C. (2014). Automated monitoring and quantitative analysis of feeding behaviour in Drosophila. Nat. Commun. 5, 4560. 10.1038/ncomms5560.

47. Davis, J.D., and Smith, G.P. (1992). Analysis of the microstructure of the rhythmic tongue movements of rats ingesting maltose and sucrose solutions. Behav. Neurosci. 106, 217–228.

48. Davie, K., Janssens, J., Koldere, D., De Waegeneer, M., Pech, U., Kreft, Ł., Aibar, S., Makhzami, S., Christiaens, V., Bravo González-Blas, C., et al. (2018). A Single-Cell Transcriptome Atlas of the Aging Drosophila Brain. Cell 174, 982–998.e20. 10.1016/j.cell.2018.05.057.

49. Dolezelova, E., Nothacker, H.-P., Civelli, O., Bryant, P.J., and Zurovec, M. (2007). A Drosophila adenosine receptor activates cAMP and calcium signaling. Insect Biochem. Mol. Biol. 37, 318–329. 10.1016/j.ibmb.2006.12.003.

50. Sayin, S., De Backer, J.-F., Siju, K.P., Wosniack, M.E., Lewis, L.P., Frisch, L.-M., Gansen, B., Schlegel, P., Edmondson-Stait, A., Sharifi, N., et al. (2019). A Neural Circuit Arbitrates between Persistence and Withdrawal in Hungry Drosophila. Neuron 104, 544–558.e6. 10.1016/j.neuron.2019.07.028.

51. McGuire, S.E., Le, P.T., Osborn, A.J., Matsumoto, K., and Davis, R.L. (2003). Spatiotemporal rescue of memory dysfunction in Drosophila. Science 302, 1765–1768. 10.1126/science.1089035.

52. Holcroft, C.E., Jackson, W.D., Lin, W.-H., Bassiri, K., Baines, R.A., and Phelan, P. (2013). Innexins Ogre and Inx2 are required in glial cells for normal postembryonic development of the *Drosophila* central nervous system. J. Cell Sci., jcs.117994. 10.1242/jcs.117994.

53. Weiss, S., Clamon, L.C., Manoim, J.E., Ormerod, K.G., Parnas, M., and Littleton, J.T. (2022). Glial ER and GAP junction mediated Ca ^2+^ waves are crucial to maintain normal brain excitability. Glia 70, 123–144. 10.1002/glia.24092.

54. de Tredern, E., Rabah, Y., Pasquer, L., Minatchy, J., Plaçais, P.-Y., and Preat, T. (2021). Glial glucose fuels the neuronal pentose phosphate pathway for long-term memory. Cell Rep. 36, 109620. 10.1016/j.celrep.2021.109620.

55. Fosque, B.F., Sun, Y., Dana, H., Yang, C.-T., Ohyama, T., Tadross, M.R., Patel, R., Zlatic, M., Kim, D.S., Ahrens, M.B., et al. (2015). Neural circuits. Labeling of active neural circuits in vivo with designed calcium integrators. Science 347, 755–760. 10.1126/science.1260922.

56. Moeyaert, B., Holt, G., Madangopal, R., Perez-Alvarez, A., Fearey, B.C., Trojanowski, N.F., Ledderose, J., Zolnik, T.A., Das, A., Patel, D., et al. (2018). Improved methods for marking active neuron populations. Nat. Commun. 9, 4440. 10.1038/s41467-018-06935-2.

57. Dus, M., Ai, M., and Suh, G.S.B. (2013). Taste-independent nutrient selection is mediated by a brain-specific Na+/solute co-transporter in Drosophila. Nat. Neurosci. 16, 526–528. 10.1038/nn.3372.

58. Plaçais, P.-Y., and Preat, T. (2013). To Favor Survival Under Food Shortage, the Brain Disables Costly Memory. Science 339, 440–442. 10.1126/science.1226018.

59. Klapoetke, N.C., Murata, Y., Kim, S.S., Pulver, S.R., Birdsey-Benson, A., Cho, Y.K., Morimoto, T.K., Chuong, A.S., Carpenter, E.J., Tian, Z., et al. (2014). Independent optical excitation of distinct neural populations. Nat. Methods 11, 338–346. 10.1038/nmeth.2836.

60. Nagel, G., Szellas, T., Huhn, W., Kateriya, S., Adeishvili, N., Berthold, P., Ollig, D., Hegemann, P., and Bamberg, E. (2003). Channelrhodopsin-2, a directly light-gated cation-selective membrane channel. Proc. Natl. Acad. Sci. 100, 13940–13945. 10.1073/pnas.1936192100.

61. Perea, G., Yang, A., Boyden, E.S., and Sur, M. (2014). Optogenetic astrocyte activation modulates response selectivity of visual cortex neurons in vivo. Nat. Commun. 5, 3262. 10.1038/ncomms4262.

62. Moreira, J.-M., Itskov, P.M., Goldschmidt, D., Baltazar, C., Steck, K., Tastekin, I., Walker, S.J., and Ribeiro, C. (2019). optoPAD, a closed-loop optogenetics system to study the circuit basis of feeding behaviors. eLife 8, e43924. 10.7554/eLife.43924.

63. Gomez, J.A., Perkins, J.M., Beaudoin, G.M., Cook, N.B., Quraishi, S.A., Szoeke, E.A., Thangamani, K., Tschumi, C.W., Wanat, M.J., Maroof, A.M., et al. (2019). Ventral tegmental area astrocytes orchestrate avoidance and approach behavior. Nat. Commun. 10, 1455. 10.1038/s41467-019-09131-y.

64. Requie, L.M., Gómez-Gonzalo, M., Speggiorin, M., Managò, F., Melone, M., Congiu, M., Chiavegato, A., Lia, A., Zonta, M., Losi, G., et al. (2022). Astrocytes mediate long-lasting synaptic regulation of ventral tegmental area dopamine neurons. Nat. Neurosci. 25, 1639–1650. 10.1038/s41593-022-01193-4.

65. Yang, J., Chen, J., Liu, Y., Chen, K.H., Baraban, J.M., and Qiu, Z. (2023). Ventral tegmental area astrocytes modulate cocaine reward by tonically releasing GABA. Neuron 111, 1104–1117.e6. 10.1016/j.neuron.2022.12.033.

66. Siju, K.P., Štih, V., Aimon, S., Gjorgjieva, J., Portugues, R., and Grunwald Kadow, I.C. (2020). Valence and State-Dependent Population Coding in Dopaminergic Neurons in the Fly Mushroom Body. Curr. Biol. 30, 2104–2115.e4. 10.1016/j.cub.2020.04.037.

67. Zolin, A., Cohn, R., Pang, R., Siliciano, A.F., Fairhall, A.L., and Ruta, V. (2021). Context-dependent representations of movement in Drosophila dopaminergic reinforcement pathways. Nat. Neurosci. 24, 1555–1566. 10.1038/s41593-021-00929-y.

68. May, C.E., Rosander, J., Gottfried, J., Dennis, E., and Dus, M. (2020). Dietary sugar inhibits satiation by decreasing the central processing of sweet taste. eLife 9, e54530. 10.7554/eLife.54530.

69. Aimon, S., Cheng, K.Y., Gjorgjieva, J., and Grunwald Kadow, I.C. (2023). Global change in brain state during spontaneous and forced walk in Drosophila is composed of combined activity patterns of different neuron classes. eLife 12, e85202. 10.7554/eLife.85202.

70. Schirmeier, S., and Klämbt, C. (2015). The Drosophila blood-brain barrier as interface between neurons and hemolymph. Mech. Dev. 138, 50–55. 10.1016/j.mod.2015.06.002.

71. McMullen, E., Hertenstein, H., Strassburger, K., Deharde, L., Brankatschk, M., and Schirmeier, S. (2023). Glycolytically impaired Drosophila glial cells fuel neural metabolism via β-oxidation. Nat. Commun. 14, 2996. 10.1038/s41467-023-38813-x.

72. Delgado, M.G., Oliva, C., López, E., Ibacache, A., Galaz, A., Delgado, R., Barros, L.F., and Sierralta, J. (2018). Chaski, a novel Drosophila lactate/pyruvate transporter required in glia cells for survival under nutritional stress. Sci. Rep. 8, 1186. 10.1038/s41598-018-19595-5.

73. Silva, B., Mantha, O.L., Schor, J., Pascual, A., Plaçais, P.-Y., Pavlowsky, A., and Preat, T. (2022). Glia fuel neurons with locally synthesized ketone bodies to sustain memory under starvation. Nat. Metab. 4, 213–224. 10.1038/s42255-022-00528-6.

74. Rabah, Y., Francés, R., Minatchy, J., Guédon, L., Desnous, C., Plaçais, P.-Y., and Preat, T. (2023). Glycolysis-derived alanine from glia fuels neuronal mitochondria for memory in Drosophila. Nat. Metab. 5, 2002–2019. 10.1038/s42255-023-00910-y.

75. Haynes, P.R., Pyfrom, E.S., Li, Y., Stein, C., Cuddapah, V.A., Jacobs, J.A., Yue, Z., and Sehgal, A. (2024). A neuron-glia lipid metabolic cycle couples daily sleep to mitochondrial homeostasis. Nat. Neurosci. 27, 666–678. 10.1038/s41593-023-01568-1.

76. Pellerin, L., and Magistretti, P.J. (1994). Glutamate uptake into astrocytes stimulates aerobic glycolysis: a mechanism coupling neuronal activity to glucose utilization. Proc. Natl. Acad. Sci. 91, 10625–10629. 10.1073/pnas.91.22.10625.

77. Theparambil, S.M., Kopach, O., Braga, A., Nizari, S., Hosford, P.S., Sagi-Kiss, V., Hadjihambi, A., Konstantinou, C., Esteras, N., Gutierrez Del Arroyo, A., et al. (2024). Adenosine signalling to astrocytes coordinates brain metabolism and function. Nature. 10.1038/s41586-024-07611-w.

78. Nortley, R., and Attwell, D. (2017). Control of brain energy supply by astrocytes. Curr. Opin. Neurobiol. 47, 80–85. 10.1016/j.conb.2017.09.012.

79. Halassa, M.M., Florian, C., Fellin, T., Munoz, J.R., Lee, S.-Y., Abel, T., Haydon, P.G., and Frank, M.G. (2009). Astrocytic Modulation of Sleep Homeostasis and Cognitive Consequences of Sleep Loss. Neuron 61, 213–219. 10.1016/j.neuron.2008.11.024.

80. Martin-Fernandez, M., Jamison, S., Robin, L.M., Zhao, Z., Martin, E.D., Aguilar, J., Benneyworth, M.A., Marsicano, G., and Araque, A. (2017). Synapse-specific astrocyte gating of amygdala-related behavior. Nat. Neurosci. 20, 1540–1548. 10.1038/nn.4649.

81. Illes, P., Burnstock, G., and Tang, Y. (2019). Astroglia-Derived ATP Modulates CNS Neuronal Circuits. Trends Neurosci. 42, 885–898. 10.1016/j.tins.2019.09.006.

82. Corkrum, M., Covelo, A., Lines, J., Bellocchio, L., Pisansky, M., Loke, K., Quintana, R., Rothwell, P.E., Lujan, R., Marsicano, G., et al. (2020). Dopamine-Evoked Synaptic Regulation in the Nucleus Accumbens Requires Astrocyte Activity. Neuron 105, 1036–1047.e5. 10.1016/j.neuron.2019.12.026.

83. Chen, A.B., Duque, M., Wang, V.M., Dhanasekar, M., Mi, X., Rymbek, A., Tocquer, L., Narayan, S., Prober, D., Yu, G., et al. (2024). Norepinephrine changes behavioral state via astroglial purinergic signaling. Preprint, 10.1101/2024.05.23.595576 10.1101/2024.05.23.595576.

84. Aimon, S., Katsuki, T., Jia, T., Grosenick, L., Broxton, M., Deisseroth, K., Sejnowski, T.J., and Greenspan, R.J. (2019). Fast near-whole-brain imaging in adult Drosophila during responses to stimuli and behavior. PLoS Biol. 17, e2006732. 10.1371/journal.pbio.2006732.

85. Zhang, Y.V., Ormerod, K.G., and Littleton, J.T. (2017). Astrocyte Ca ^2+^ Influx Negatively Regulates Neuronal Activity. eneuro 4, ENEURO.0340-16.2017. 10.1523/ENEURO.0340-16.2017.

86. Otto, N., Marelja, Z., Schoofs, A., Kranenburg, H., Bittern, J., Yildirim, K., Berh, D., Bethke, M., Thomas, S., Rode, S., et al. (2018). The sulfite oxidase Shopper controls neuronal activity by regulating glutamate homeostasis in Drosophila ensheathing glia. Nat. Commun. 9, 3514. 10.1038/s41467-018-05645-z.

87. Liu, H., Zhou, B., Yan, W., Lei, Z., Zhao, X., Zhang, K., and Guo, A. (2014). Astrocyte-like glial cells physiologically regulate olfactory processing through the modification of ORN - PN synaptic strength in *D rosophila*. Eur. J. Neurosci. 40, 2744–2754. 10.1111/ejn.12646.

88. Coman, C., Solari, F.A., Hentschel, A., Sickmann, A., Zahedi, R.P., and Ahrends, R. (2016). Simultaneous Metabolite, Protein, Lipid Extraction (SIMPLEX): A Combinatorial Multimolecular Omics Approach for Systems Biology. Mol. Cell. Proteomics 15, 1435–1466. 10.1074/mcp.M115.053702.

89. El Abiead, Y., Bueschl, C., Panzenboeck, L., Wang, M., Doppler, M., Seidl, B., Zanghellini, J., Dorrestein, P.C., and Koellensperger, G. (2022). Heterogeneous multimeric metabolite ion species observed in LC-MS based metabolomics data sets. Anal. Chim. Acta 1229, 340352. 10.1016/j.aca.2022.340352.

90. Hildebrandt, A., Bickmeyer, I., and Kühnlein, R.P. (2011). Reliable Drosophila Body Fat Quantification by a Coupled Colorimetric Assay. PLoS ONE 6, e23796. 10.1371/journal.pone.0023796.

91. Xu, Y., Borcherding, A.F., Heier, C., Tian, G., Roeder, T., and Kühnlein, R.P. (2019). Chronic dysfunction of Stromal interaction molecule by pulsed RNAi induction in fat tissue impairs organismal energy homeostasis in Drosophila. Sci. Rep. 9, 6989. 10.1038/s41598-019-43327-y.

92. Quarta, C., Fisette, A., Xu, Y., Colldén, G., Legutko, B., Tseng, Y.-T., Reim, A., Wierer, M., De Rosa, M.C., Klaus, V., et al. (2019). Functional identity of hypothalamic melanocortin neurons depends on Tbx3. Nat. Metab. 1, 222–235. 10.1038/s42255-018-0028-1.

